# Grayscale 4D Biomaterial Customization at High Resolution and Scale

**DOI:** 10.1101/2024.01.31.578280

**Authors:** Ivan Batalov, Jeremy R. Filteau, Ryan M. Francis, Georg Jaindl, Luke Orr, Teresa L. Rapp, Shiyu Yang, Jordan A. Filteau, Weisi Xie, Ross C. Bretherton, Adam K. Glaser, Jonathan T.C. Liu, Kelly R. Stevens, Cole A. DeForest

## Abstract

Hydrogel biomaterials have proven indispensable for three-dimensional (3D) cell culture but have fallen short in replicating the innate physiochemical complexity of real tissue. Though traditional photolithography affords localized material manipulation, robust methods that govern when, where, and to what extent such phototailoring occurs throughout materials would be profoundly enabling towards fabricating more-realistic 3D tissue constructs. Here, we introduce “grayscale image z-stack-guided multiphoton optical-lithography” (GIZMO) as a generalizable and intuitive strategy to rapidly photomodulate materials in full 3D non-binary patterns at submicron resolutions spanning large volumes (>mm^3^). Highlighting its versatility, we employ GIZMO to variably photopattern biomolecule release from, protein immobilization to, and degradation within hydrogels based on biologically derived or synthetic grayscale image stacks with unprecedented complexity. We anticipate that GIZMO will enable new opportunities to probe and manipulate cell fates, as well as to engineer complex functional tissue.

## Main text

Over the past three decades, hydrogels have gained unrivaled popularity as biomaterial constructs for three-dimensional (3D) cell culture and tissue engineering (*1–4*). While hydrogels provide a culture environment that recapitulates many critical aspects of the cellular niche (e.g., high water content, tissue-like viscoelasticity), modern engineered systems largely fall short in mimicking the spatiotemporal complexity and dynamics of native tissue. Though recent advances in bioprinting now permit the specification of hydrogel geometries and compositions at macroscopic scales (*5–8*), leading strategies are limited in their geometric and structural complexity (typical minimum feature size >100 µm) and are often static (i.e., once formed, they cannot be readily altered) (*9–16*). New fabrication methods are needed to replicate the dynamic biochemical/biophysical complexity of tissues from macroscopic (mm) to cellular scales (i.e., sub- or single micron).

Complementing conventional bioprinting, photochemical methods offer a route to cytocompatibly customize four-dimensional (4D) biomaterial properties at resolutions dictated theoretically by the diffraction limit of light (typically submicron). Phototunable hydrogels have been created that stiffen, soften, or undergo cyclic mechanical alterations, as well as to immobilize, release, or reversibly bind biomolecules upon optical stimulation (*17–20*). Precise modulation of such materials is enabled through techniques that control when and where photons are directed onto specific sample subvolumes (*17*, *21*, *22*). In conventional photolithography, projected light extends throughout the gel thickness in 2D geometric patterns specified using photomasks or projection systems. Although these methods afford scalable single micron-resolved patterning across large biomaterial surfaces (*23–25*), full 3D spatial control is not possible. In multiphoton-based laser-scanning lithography, modification is confined to the focal point of a femtosecond-pulsed laser; programmed scanning of that focal point in space enables intricate 3D patterning within photoresponsive materials at high resolutions (sub-micron in the xy, single-micron in z), but traditionally only in simple geometries with binary material/chemical properties that span only very small volumes (*26–31*). The utility of such discretely modified biomaterials – those whose on/off patterns may have 3D complexity but are inherently binary – towards engineering biology’s native complexity is severely limited.

Taking advantage of the light dosage-dependence of many photochemistries, some lithographic techniques have been developed that enable graded substrate modulation. Variably positioned and non-binary patterned photomasks (*32*, *33*), digital micromirror devices (*34*), and sequential laser scanning (*35*) have successfully exploited such dosed response for heterogeneous gel customization. However, no existing technique can independently vary photomodification at different z-positions within the gels’ thickness, representing a major roadblock to controlling the extent of functionalization throughout large-scale biomaterials at subcellular resolutions.

Here, we introduce a generalizable strategy to modulate biomaterials in full 3D non-binary patterns with subcellular resolutions across mm^3^ scales. Using grayscale image z-stacks to specify laser power during continuous raster scanning on a voxel-by-voxel basis, we photocustomize hydrogel properties dynamically and at high resolution. As light intensity is varied positionally on the fly, this method – referred to as “grayscale image z-stack-guided multiphoton optical-lithography” (GIZMO) – is performed rapidly and at speeds independent of pattern complexity (**Figure 1**). Highlighting the versatility of the approach and its immediate compatibility with state-of-the-art bioorthogonal photochemistries, we employ GIZMO to photopattern biomolecule release from, protein immobilization to, and degradation within hydrogel biomaterials, infusing newfound physiological complexity into synthetic cell culture platforms.

**Fig. 1.**
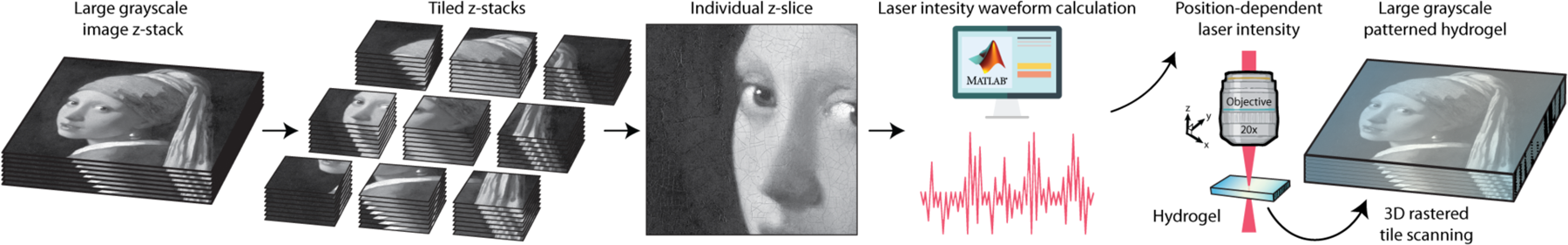
Biomaterial patterning through grayscale image z-stack-guided multiphoton optical-lithography (GIZMO). Multiphoton laser-scanning lithographic patterning enables high precision biomaterial customization in 3D, whereby photochemical reactions are confined to the focal point of a femtosecond-pulsed laser. Modulation of laser power during raster scanning as positionally specified on a voxel-by-voxel basis by the corresponding tiled z-stack location’s grayscale intensity yields patterned 3D biomaterials through GIZMO. Since inputted image stacks can be generated from 3D models (e.g., AutoCAD, STL) or obtained directly through many common imaging techniques (e.g., confocal, light sheet, two-photon microscopy), this approach affords precise control over a wide range of multiphoton-compatible chemistries following user-specified and/or biologically defined patterns.

### Grayscale image z-stack-guided multiphoton optical-lithography (GIZMO)

To ensure that gel photopatterning is robust, flexible, and intuitive, we elected to guide multiphoton-based laser-scanning lithographic processing using grayscale image z-stack templates. Such image stacks are readily acquired from native biological specimens (e.g., confocal, multiphoton, and light-sheet micrographs), exported from 3D computer-aided-design files (e.g., CAD), computationally generated, and/or obtained from repositories. Z-stacks can be processed and manipulated (e.g., resized, cropped, combined, superimposed, deconvolved, inverted, retouched) via countless commercial software and freeware packages in a manner that is inherently familiar to bioscientists, affording near-limitless pattern customization. Moreover, since they can be visualized in 3D, image stacks naturally impart a “what you see is what you get” approach to digital fabrication that is widely considered powerful and convenient.

To control the degree of photopatterning at given gel subvolumes on a voxel-by-voxel basis, we exploit individual grayscale pixel intensities to govern laser power and local light dosage in real time. This is accomplished during raster scanning through use of an acousto-optic modulator (AOM), an in-line device that can continuously vary laser output with an electrical drive signal. Using custom MATLAB code, we calculate and load electrical waveform patterns to the AOM that match its 3D outputted power profile to that of the inputted z-stack voxel intensities. Critically, since AOMs provide exceptionally high contrast (>1000:1) and operate at speeds (<1 μs full rise/fall time) faster than typical pixel dwell times used during raster scanning, such an approach can theoretically afford patterned biomaterials with properties specified non-discretely on a voxel-by-voxel basis.

We initially developed GIZMO to operate within FluoView, a proprietary software compatible only with Olympus-branded multiphoton microscopes (**Supplementary Method 1**). To ensure that GIZMO was implemented in as broadly accessible form as possible, we subsequently identified ScanImage® (Vidrio Technologies, MBF Bioscience) as a particularly powerful software package for controlling laser scanning microscopes (*36*). ScanImage® interfaces with and runs effectively on many brands of multiphoton microscopes, including those manufactured by Scientifica, Sutter, Thorlabs, and others. Built upon MATLAB, it is intrinsically open source and scriptable, permitting automated image processing that is not accessible via alternative commercial options. Taking advantage of these features, we coded GIZMO into the ScanImage® platform. After prompting the user to identify serially stacked grayscale image files and patterning parameters (e.g., laser wavelength, maximal laser intensity, scan speed), the software automatically calculates and loads electrical waveform patterns to the AOM before raster scanning the sample in the image-defined pattern. The process can be made to run exceptionally fast; using a Thorlabs Bergamo II microscope equipped with a Coherent Chameleon Discovery NX TPC femtosecond-pulsed laser with 8 kHz resonant scanning and a piezo objective “fast-z” scanner, we have obtained grayscale patterning rates that approach 30 z-frames per second (typical frame size = 360 µm x 360 µm, z-spacing = 2 µm), providing a clear path to large-scale biomaterial customization on timescales of minutes. To facilitate widespread adoption of these techniques and in partnership with Vidrio Technologies, GIZMO is now included within the current distribution of ScanImage® as the “Print3D” module.

### 3D Photopatterned Biomolecule Release from Gels via GIZMO

Following chemical strategies that have found previous success at controlling cell proliferation, differentiation, and intracellular signaling in 3D and at sub-cellular resolutions (*37–41*), we first attempted to apply GIZMO to guide biomolecule photorelease from polymeric hydrogels in well-defined grayscale patterns (**Figure 2A**). Towards this goal, we linked proteins homogenously monotagged with a photoreleasable *ortho*-nitrobenzyl ester (*o*NB) moiety to hydrogels formed through a step-growth strain-promoted azide-alkyne cycloaddition (*28*) between poly(ethylene glycol) tetra-bicyclononyne (PEG-tetraBCN, M_n_ ≈ 20 kDa, 4 mM) and a linear PEG diazide (N_3_-PEG-N_3_, M_n_ ≈ 3.5 kDa, 8 mM) in phosphate-buffered saline (pH 7.4) (**Supplementary Method S2-S8**). Upon two-photon stimulation (λ = 740 nm), the *o*NB linker undergoes irreversible cleavage to form both nitroso- and carboxylic acid-containing byproducts, triggering concomitant release of the previously immobilized species from the otherwise photostable gel (**Figure 2B**). Modification of mCherry’s C-terminus with a polyglycine-*o*NB-azide peptide [H-GGGGDDK(*o*NB-N_3_)-NH_2_] via a sortase-tag enhanced protein ligation (STEPL) (*40*, *42*) yielded a photoreleasable species (mCherry-*o*NB-N_3_) whose relative gel-bound concentration could be quantified fluorescently shortly after photopatterning, providing a direct and rapid readout of pattern fidelity. Uniquely enabled through site-specific chemoenzymatic modification (*43*), we anticipated that the monolabeled photoreleasable protein would be highly bioactive and exhibit well-behaved photokinetics.

**Fig. 2.**
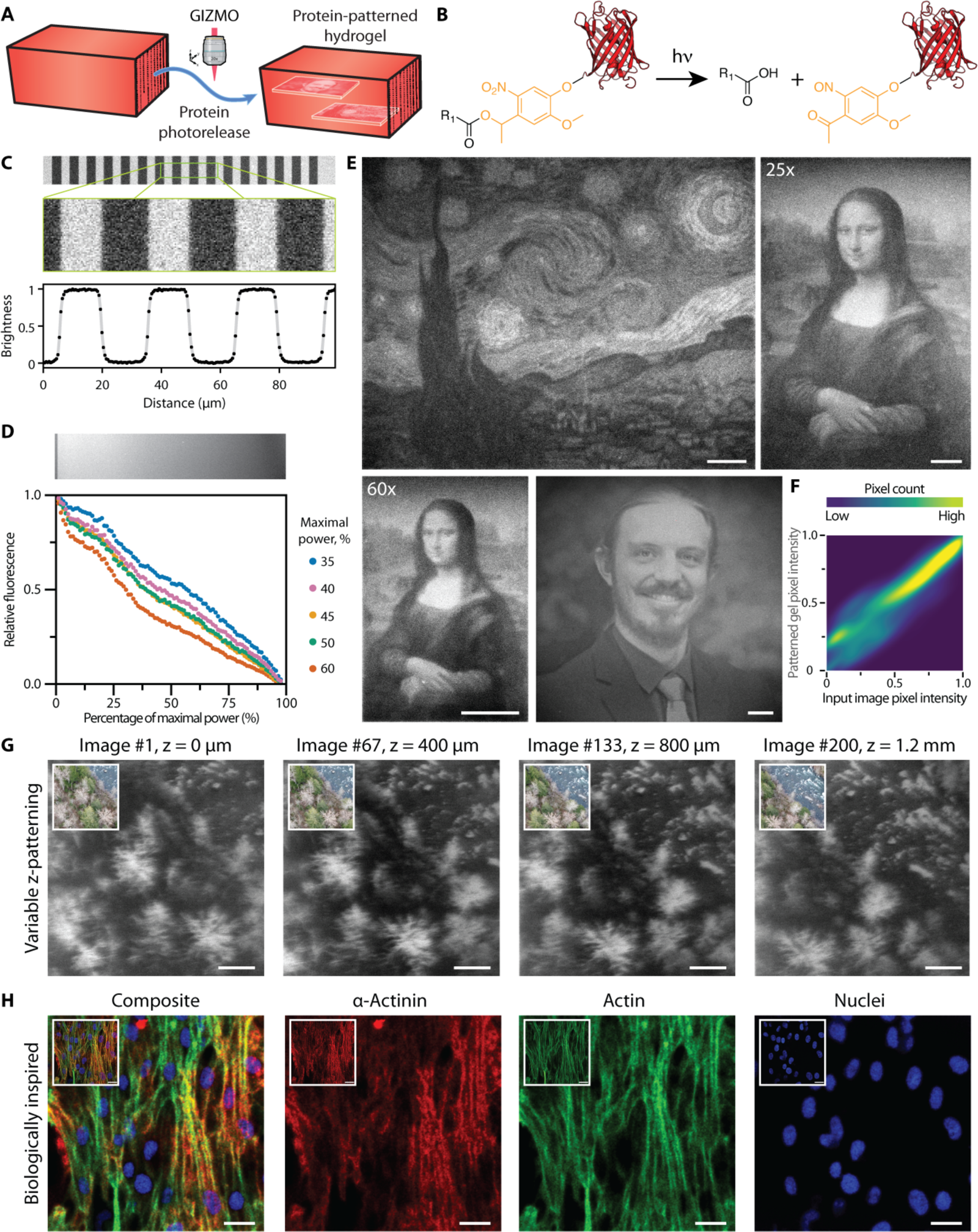
Photopatterned biomolecule release from biomaterials using GIZMO. (**A**) Site-specifically modified proteins tethered to hydrogel biomaterials through a photodegradable linker are readily photoreleased via GIZMO. (**B**). The *o*NB moiety C-terminally installed between the mCherry protein and the material (R_1_) undergoes photoscission in response to focused laser light (λ = 740 nm), resulting in protein release and its diffusive removal from the gel. (**C**) Using an interspaced line binary pattern to guide protein release, a sharp transition between areas fully functionalized and those in which protein has been completely photoreleased was achieved, as revealed by fluorescent microscopy. (**D**) Linear gradient exposure patterns (470 µm wide, varied from 0 to maximum laser intensity) conducted at different maximum laser power (35 – 60%) were used to correlate laser power with extent of mCherry protein photorelease. Representative gradient corresponds to maximum power of 40%. (**E**) GIZMO patterning of mCherry photorelease from PEG hydrogels in arbitrary patterns, including van Gogh’s “The Starry Night”, da Vinci’s “Mona Lisa”, and the photograph of a current-day biomaterials scientist. (**F**) Direct comparison of the pixel intensities from the “Mona Lisa”-patterned gel with the corresponding input image showcased high parity. (**G**) Representative slices of GIZMO-patterned grayscale z-stack consisting of 200 distinct planar images comprising the biomolecule “flipbook” (300 µm x 300 µm x 1.2 mm total size), derived from drone footage acquired near Kanaskat-Palmer Washington State Park. (**H**) Cardiomyocytes stained for α-actinin (red), actin (green), and nuclei (blue) were used as the basis for biologically defined GIZMO patterning of mCherry protein release from a gel. Patterned proteins are shown with false coloring to match that of the corresponding original fluorescent channel. Each channel was patterned within separate locations in the gel prior to superposition. For C-E and G, bound species is shown in white; released regions appear black. Inset images correspond to template images. Scale bars: 50 µm in C-F, 20 µm in G.

To probe the x-dimensional patterning resolution of GIZMO for protein photorelease, whereby the laser focus translates linearly in the x dimension during raster scanning, we first released mCherry from z locations well below the gel surface following an interspaced line binary image pattern (line width = 15 µm, line spacing = 15 µm). Using conditions standard for multiphoton imaging (pixel size = 0.5 µm/px, pixel dwell time = 2 µs/px, 11 = 740 nm, power = 40% of maximum, repeats = 32, 25× objective), we observed a sharp transition between gel regions fully functionalized with protein and those where it had been completely released (**Figure 2C**). Fluorescent analysis further revealed a transition resolution distance of approximately 2 – 3 pixels (90% = 2 px ∼ 1.1 µm, 95% = 3 px ∼ 1.4 µm), as imaged. Since the time required for the AOM to tune the laser intensity is proportional to power change, which is maximal in the case of a binary pattern, this was viewed as the “worst case” x resolution for the stated conditions.

We next employed GIZMO for grayscale patterned protein release from hydrogels, first utilizing an image file uniform in the y dimension but graded linearly across all pixel intensities (0–100) over ∼470 µm in x. Varying the maximal laser power from 35 – 60% but holding all other scan parameters constant, the expected gradients were formed (**Figure 2D**). Though *o*NB photocleavage follows non-linear kinetics, we observed that a near-linear graded response was achieved when a maximal laser power of 40% was employed. Utilizing this laser power setting, grayscale images could be faithfully reproduced in hydrogels without the use of a predetermined lookup table correlating laser power to the extent of protein release. In this manner, we exploited GIZMO (25× objective) to reproduce Vincent van Gogh’s “The Starry Night”, Leonardo da Vinci’s “Mona Lisa”, and the photograph of a current-day biomaterial scientist in protein-functionalized gels, each with excellent fidelity achieved over single seconds (**Figure 2E**).

Utilization of a high-magnification objective (60×) enabled even finer grained features to be formed. These GIZMO patterns are created with significantly higher grayscale depth and with a method that is at least two orders of magnitude faster than serial scanning (**Supplementary Figure S1**). Parity plots comparing the pixel intensity of the inputted image with that from the patterned gels demonstrated high similarity and effective patterning (**Figure 2F, Supplementary Fig. S2**).

Taking GIZMO to the next dimension, we patterned photorelease of a fluorescent bioactive peptide [N_3_-*o*NB-GRGDSK(AF488)-NH_2_, **Supplementary Method S9**] from gels using a grayscale z-stack consisting of 200 distinct planar images using previously optimized conditions. The image stack was generated from author-acquired drone footage of the forested Green River near the Kanaskat-Palmer Washington State Park, mapping the video’s time dimension onto z-stack position. Using a z-spacing of 6 µm, we patterned (25× objective) this grayscale image stack within our gel, resulting in a 300 µm x 300 µm x 1.2 mm digitally fabricated biomolecule “flipbook” (**Figure 2G, Supplementary Movie S1**). Pattern fidelity remained high, independent of sample location, affording a patterned hydrogel with unprecedented 3D complexity and providing a critical step towards recapitulating native tissue’s diverse biochemical makeup.

Seeking to demonstrate that GIZMO enables biologically directed microfabrication, we patterned protein release from gels (60× objective) following fluorescent confocal micrographs of immunostained cardiomyocytes cultured on glass. Localizing mCherry release to inverted patterns of cardiomyocyte cytoskeletal α-actinin, actin skeletal muscle, and nuclei fluorescent channels, each at a different gel position, demonstrated 1:1 biomaterial customization on a native biological scale (**Figure 2H**). Moving forward, we anticipate that such bioinspired fabrication will provide a uniquely powerful tool to study and recreate functional tissues.

### 3D Protein Photoimmobilization within Biomaterials via GIZMO

Building on success in grayscale protein photorelease from biomaterials, we turned our attention towards grayscale patterned immobilization of proteins throughout hydrogels. Though biochemical patterning of hydrogels has proven effective in spatially guiding cell fate, multiphoton-enabled protein tethering within gels has been only sparsely explored (*30*, *35*, *38*– *40*, *44–46*). Prior studies have yielded only binary patterns with limited 3D complexity, with the one notable exception reported by the Zenobi-Wong group where a small grayscale protein pattern invariant in the z dimension was demonstrated (*35*). Towards filling this technological gap, we sought to apply GIZMO in conjunction with a bioorthogonal oxime photoligation (*38*, *40*, *44*, *46*) to immobilize site-specifically modified proteins within polymeric hydrogels in well-defined non-binary patterns across cubic millimeter volumes (**Figure 3A**).

**Fig. 3.**
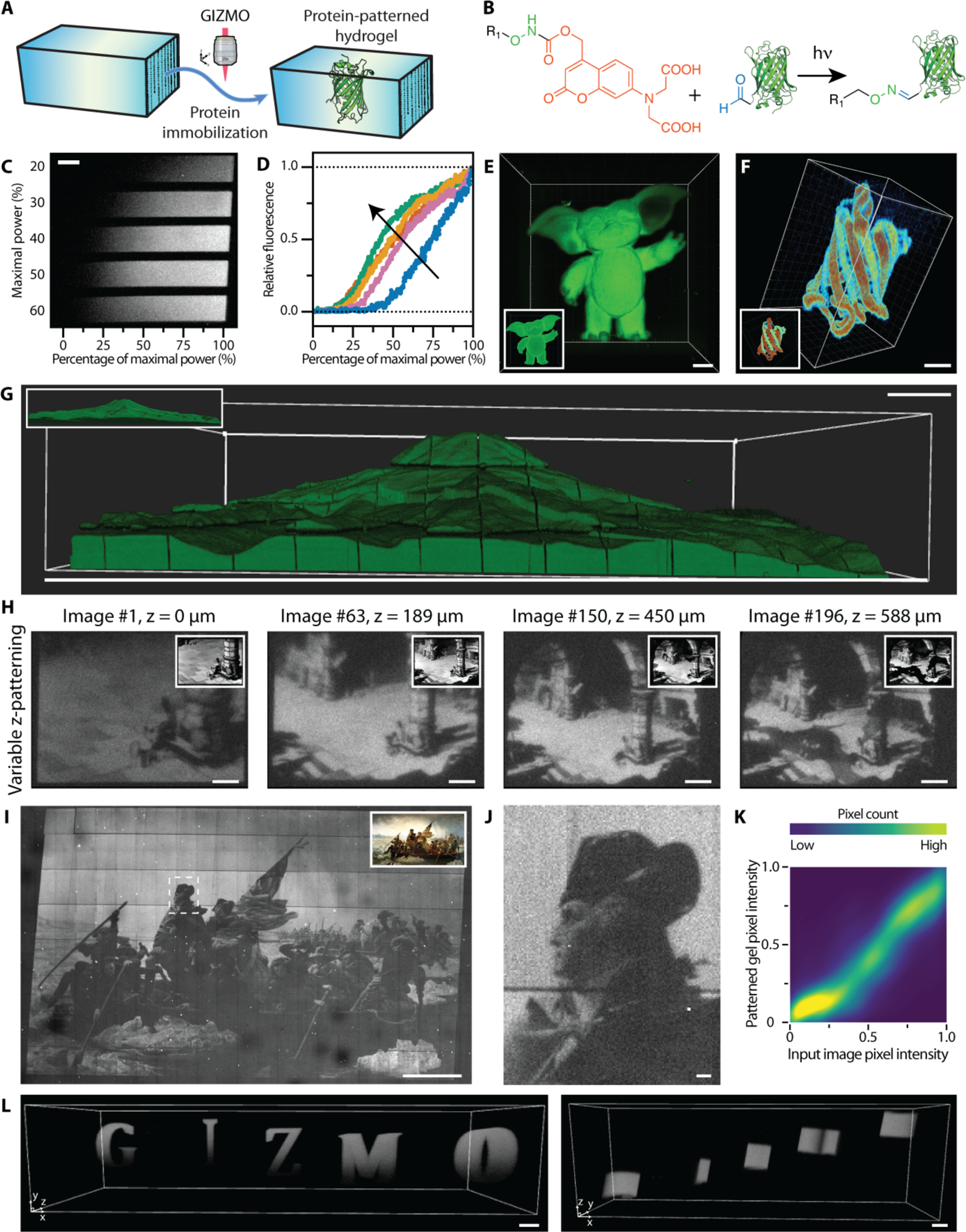
Photopatterned protein immobilization within hydrogels via GIZMO. (**A**) Site-specifically modified proteins are photochemically tethered at specific regions within biomaterials by GIZMO. (**B**) A photomediated oxime chemistry permits aldehyde-modified proteins to be covalently tethered C-terminally within the gel in response to pulsed laser light (λ = 740 nm). (**C-D**) Linear gradient exposure patterns (varied from 0 to maximum laser intensity) at different maximal laser powers (20 – 60%) were used to relate protein immobilization with patterning conditions. Arrow denotes increasing maximal power (20, 30, 40, 50, 60%). **E**) 3D patterned immobilization of EGFP-CHO is achieved within the gel in the shape of a lovable gremlin. (**F**) Gels were functionalized with EGFP-CHO in a 3D geometry as defined by the protein’s crystal structure, with alternating strands corresponding to different levels of modification (color map from red to green with increased protein functionalization). (**G**) Tiled binary patterned immobilization of mGL-CHO in the form of Mount Rainier (8 mm x 8 mm x 2 mm, approximate 1:3,000,000 scale). (**H**) Representative slices of GIZMO-patterned grayscale z-stack consisting of 200 distinct planar images comprising the mGL-CHO “flipbook” (400 µm x 300 µm x 600 µm total size), in the form of Walt Disney’s short film “The Mad Doctor”. (**I-K**) 2D tiled GIZMO photoimmobilization of mGL-CHO in the form of Leutze’s “Washington Crossing the Delaware”. Pixel intensity comparisons showcase high parity between input image and patterned gels. (**L**) Tiled GIZMO gel patterning for 3D letters “GIZMO”. Individual letters were patterned with linearly graded concentration in y, as well as offset positions in z. For C, H-J, and L, immobilized species is shown in white; unfunctionalized regions appear black. Inset images correspond to reference images and 3D models that guided GIZMO. Scale bars: 50 µm in C-F and J, 100 µm in H and L, 500 µm in G and I.

Photodecoratable hydrogels were formed via SPAAC step polymerization of PEG-tetraBCN (M_n_ ≈ 20 kDa, 4 mM), tri(ethylene glycol) diazide (8 mM), and a heterobifunctional tetra(ethylene glycol) bearing both a BCN and a 7-dicarboxymethylaminocoumarin (DCMAC)-photocaged alkoxyamine (DCMAC-HNO-TEG-BCN, 100 µM, **Supplementary Methods S10-11**). Though not previously utilized for photomediated oxime ligation, we anticipated that DCMAC’s high two-photon sensitivity (*35*, *47*) would prove useful for GIZMO-based gel customization. Upon two-photon stimulation (λ = 740 nm), DCMAC is cleaved, liberating a reactive alkoxyamine that undergoes localized oxime ligation with gel-swollen aromatic aldehyde-modified proteins (**Figure 3B**). Following diffusive removal of unbound proteins, the gel remains covalently functionalized in light-exposed regions. Chemoenzymatic modification of the C-termini of both Enhanced Green Fluorescent Protein (EGFP) and mGreenLantern (mGL (*48*)) with a polyglycine-aldehyde peptide [H-GGGGDDK(CHO)-NH_2_, **Supplementary Methods S3-4 and S6-7**] via STEPL yielded homogenously monotagged and photopatternable species (EGFP-CHO, mGL-CHO) whose fluorescence could be used to visualize and assess patterning success.

After determining optimal laser conditions for grayscale protein phototethering (pixel size = 0.78 µm/px, pixel dwell time = 88 ns, 11 = 770 nm, power = 80% of maximum, repeats = 10, 25× objective) (**Figures 3C-D**), we fed GIZMO sliced 3D model inputs (STL files) to immobilize EGFP-CHO locally within gels in the 3D shape of the Seattle Space Needle, a lovable gremlin, and a two-toned version of the protein’s crystal structure (Protein Data Bank ID: 2Y0G (*49*)) (**Figures 3E-F, Supplementary Figure S3**). Taking advantage of DCMAC’s high two-photon sensitivity, we employed tiled GIZMO to tether mGL-CHO within hydrogels matching the shape of Mount Rainier (**Figure 3G**). Notably, patterning time of this large structure (8 mm x 8 mm x 2 mm, 130 µL, 1:3,000,000 scale) was complete in just under 7 hours. Using an inputted image stack (200 images) from Walt Disney’s short film “The Mad Doctor” (public domain), a mGL-based “flipbook” (900 µm x 700 µm x 600 µm total size) was also patterned (**Figure 3H**, **Supplementary Movie S2**). Tiled implementation of GIZMO enabled grayscale protein immobilization across large scales in 3D and with excellent pattern fidelity, evidenced through recreation of Leutze’s “Washington Crossing the Delaware” (3.8 mm x 2.4 mm) and z-graded letters corresponding to our technique’s acronym (2.7 mm x 0.8 mm x 0.7 mm) (**Figures 3I-L**). We anticipate that such 3D grayscale biochemical modulation will prove useful in probing and directing cell fate, particularly when the image stacks driving GIZMO are derived directly from living samples.

### Precisely Controlled 3D Hydrogel Degradation via GIZMO

Building on success in customizing grayscale biomaterial biochemistry, we sought to further highlight the versatility of GIZMO by using it to create well-defined synthetic microvasculature within hydrogels (**Figure 4A**). Though small multiphoton-patterned microvoids within gels have found substantial utility in controlling multicellular geometry and outgrowth (*50–53*), creating perfusable and cellularized networks (*54–57*), and in investigating biological mechanisms underpinning hematologic diseases (*58*), prior efforts have been substantially limited in their 3D complexity and size. We aimed to demonstrate that GIZMO could be applied in light-mediated subtractive manufacturing to readily create synthetic microvasculature with capillary-sized features whose makeup is physiologically informed.

**Fig. 4.**
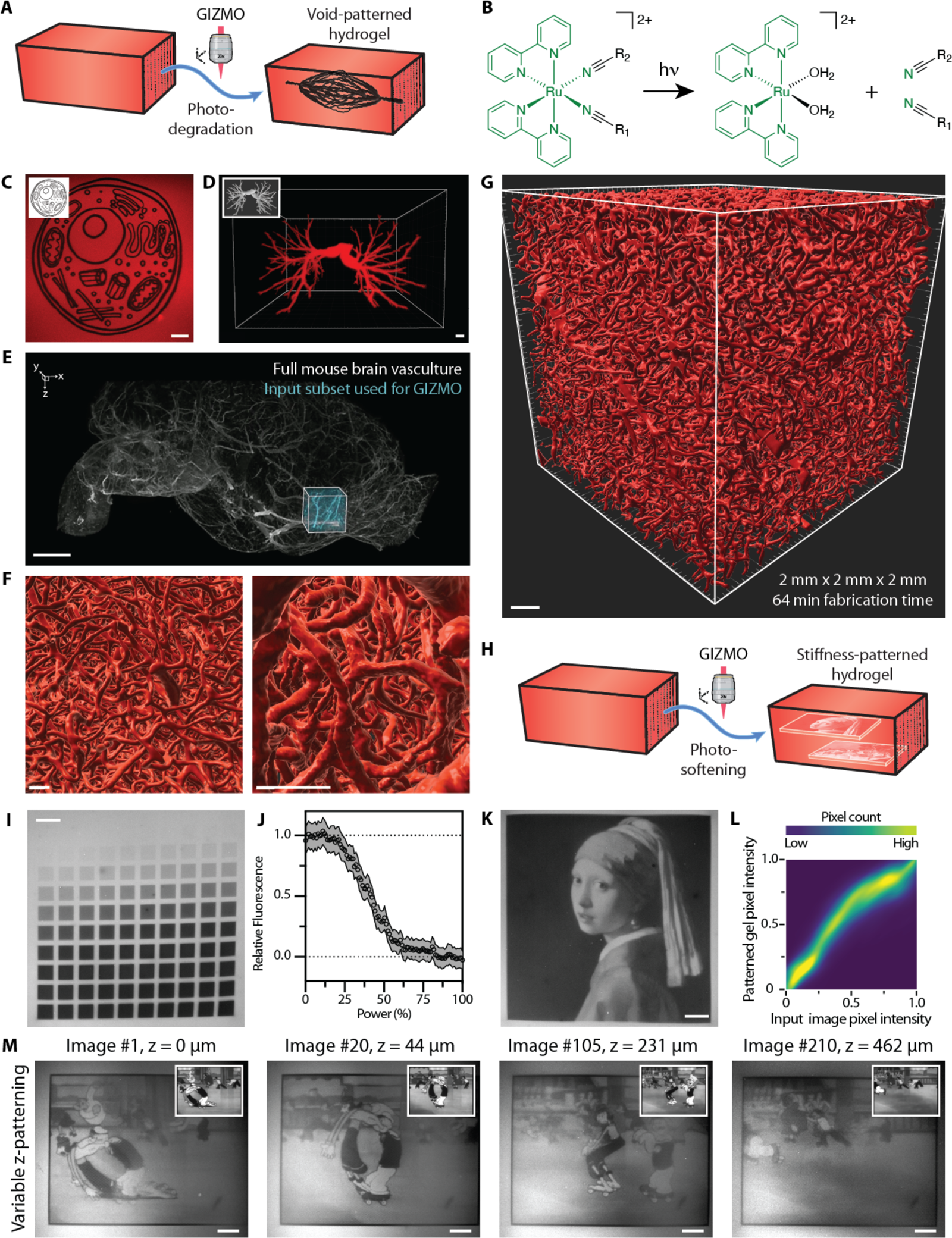
Patterned biomaterial photodegradation through GIZMO. (**A**) Patterned 3D voids are readily etched within photodegradable hydrogels via GIZMO. (**B**) The Rubpy crosslinkers within the gel backbone (R_1_ and R_2_) photolyze under focused laser light (λ = 750 – 850 nm) irradiation, locally inducing reverse gelation and material dissolution. (**C**) 2D control over material degradation is achieved within 3D gels in arbitrary patterns. (**D**) 3D patterning of an anatomical model of human pulmonary arteries is achieved through GIZMO-driven hydrogel photodegradation, showcased with a 3D color-inverted rendering. (**E**) Entire intact mouse brain with fluorescently stained vasculature (grey, anti-aSMA staining; teal, input subset used for GIZMO), obtained by open-top light-sheet microscopy following CUBIC clearing, and subset used for vessel patterning (cyan). (**F-G**) Surface renderings of tiled GIZMO-photodegraded voids throughout gel at various magnifications, following native microvasculature pattern in E. (**H**) Partial material photodegradation by GIZMO affords patterned material softening. (**I-J**) Gels exposed to a gridded array of varying power (0% – 100% relative to maximum, moving from left-to-right, top-to-bottom). Following fluorescent staining, extent of gel photodegradation was assessed through loss of fluorescence, and (J) quantified with bands reflecting error about the mean for n=3. (**K**) 2D GIZMO-patterned photosoftening of gel following Vermeer’s “Girl with a Pearl Earring”. (**L**) High parity on a pixel-by-pixel basis between inputted image and patterned gel softening was obtained. (**M**) Representative slices of GIZMO-patterned softened hydrogel “flipbook” consisting of 300 planar images (450 µm x 350 µm x 650 µm total size), in the form of the Popeye short film “A Date to Skate”. For C and I-M, intact gel is shown in white or red, while photodegraded regions appear black; grayscale values in between correspond to partially degraded materials of intermediate stiffnesses. Inset images correspond to reference images and 3D models that guided GIZMO. Scale bars: 50 µm in C and I-M, 10 µm in D, 2 mm in E, 50 µm in F, 200 µm in G.

Towards this goal, photodegradable hydrogels were formed via SPAAC step polymerization of PEG-tetraBCN (M_n_ ≈ 20 kDa, 4.05 mM) and a recently reported bis-azide modified photodegradable ruthenium (Ru) polypyridyl complex-based crosslinker [(Ru(2,2’-bipyridine)_2_-(4-azidobutanenitrile)_2_, Rubpy (*59*), 8 mM] (**Supplementary Method S12**). After gel formation, two-photon absorption (11 = 750 – 850 nm) and triggered crosslinker photoscission permits direct hydrogel restructuring through localized material dissolution (**Figure 4B**). Taking advantage of the slight stoichiometric excess of BCN groups included during gel formulation, photopatterned gels were stained with a fluorescent azide dye (Cy5-N_3_, 1.25 µM). After gel patterning and fluorescent labeling, we imaged samples using fluorescence confocal microscopy and quantified the extent of degradation within and immediately surrounding each of the illuminated regions (here, a lack of fluorescence was taken to indicate complete photodegradation). This label-after-photopatterning approach enables full confidence that desired gel regions were degraded as intended (**Supplementary Figure S4**).

We first identified optimal laser scanning parameters enabling complete photodegradation, defined as those that afforded the fastest (e.g., minimal scan repeats) patterning with minimal bleed in the z dimension. To our delight, Rubpy proved exceptionally sensitive to two-photon-mediated cleavage, enabling complete and localized material degradation following a single resonance-scanned (8 kHz) exposure at full power (pixel size = 0.70 µm/px, pixel dwell time = 88 ns/px, λ = 750 nm, repeat = 1, 25× objective). Using these parameters, GIZMO was employed to photodegrade 2D and patent 3D structures into gels, initially within one imaging window (**Figures 4C-D, Supplementary Figure S5**).

Towards generating large-scale, native tissue-derived vasculature networks, we cleared, stained (anti-α-Smooth Muscle Actin), and 3D imaged the vascular network within whole mouse brain samples using hybrid open-top light-sheet microscopy (*60*) (**Supplementary Method S13**). The entire intact brain was imaged at a rate of ∼350 mm^3^ hr^-1^ with near-isotropic resolution (*xyz* resolution of 4.41 ± 0.83, 4.09 ± 1.07, and 5.48 ± 1.08 μm). We next developed a custom Python-based workflow for 3D vessel segmentation, which we used to generate a high-resolution 3D model for a 2 mm x 2 mm x 2 mm vasculature subvolume (**Figure 4E**, **Supplementary Method S14**). This model was further transformed into a tiled image stack used to drive GIZMO-based gel photodegradation. Owing the Rubpy’s remarkable two-photon sensitivity, the entire GIZMO process took just 64 min for the 8 mm^3^ structure. Furthermore, results highlight that GIZMO’s underlying AOM was able to effectively modulate laser power with high x-resolution, even under such rapid laser scanning. Upon fluorescent labeling and visualization (color inversion-based surface renderings employed for 3D visualization), patent capillary-sized channels that were faithfully indistinguishable from the inputted model were observed with diameters spanning single to hundreds of micrometers over cubic millimeter volumes (**Figure 4F-G, Supplementary Movies S3-4**). These patterns extend well beyond the current state of the art for engineered vasculature networks in complexity, size, and resolution.

Finally, recognizing the importance of networks mechanics on cell function (*61*), we sought to exploit partial material photodegradation to manufacture hydrogels with locally varied grayscale mechanics (**Figure 4H**). Towards this, we determined the grayscale photodegradation response to varied laser power, holding all other scan parameters constant (pixel size = 0. 44 µm/px, pixel dwell time = 3200 ns/px, 11 = 850 nm, repeats = 1, 25× objective) (**Figures 4I-J**). Stiffness-patterned gels were then created in the form of Vermeer’s “Girl with a Pearl Earring” and many others, each with high pattern fidelity (**Figure 4K-L, Supplementary Figure S6**). Using similar scanning parameters, we created a mechanically variable “flipbook” following inputted frames from the Popeye short film “A Date to Skate” (public domain) consisting of 300 planar images (450 µm x 350 µm x 650 µm total size) (**Figure 4M**, **Supplementary Movie S5**). We anticipate that the ability to customize network stiffness at subcellular resolutions and in 3D will prove useful for mechanobiology, just as the engineered voids will aid in building vascularized tissues, fabricating next-generation microfluidic devices, and other applications that require precise microstructural control.

## Conclusion

In this manuscript, we introduced GIZMO as a generalizable strategy to rapidly photocustomize mm^3^-scale materials at submicron resolutions and with 4D grayscale control. We have demonstrated GIZMO’s specific use in patterning non-discrete protein release from, biomacromolecule immobilization to, and degradation within polymeric hydrogels at unprecedented complexity and scale. The method is largely photochemically agnostic and is expected to be compatible with the ever-expanding library of multiphoton-sensitive chemistries. To facilitate widespread and rapid adoption by the community, we have prioritized open-source code development that ensures compatibility with several leading commercial as well as homebuilt multiphoton microscopes.

Modern biological research increasingly suggests a dense interconnectedness between biological structure, proximity, and function. We anticipate that the threads of this network will be increasingly revealed by flexible and high-throughput tools that can probe, direct, and eventually replicate biological function in heterogenous tissue culture. Beyond biological research, engineering challenges such as personalized medicine, drug discovery, and biological data storage will benefit from robust image-guided biomaterial fabrication. GIZMO represents a major step toward these challenges as the first platform for rapid, high-resolution, and grayscale photomodulation of biomaterials in 3D.

## Acknowledgements

We thank J. Davis, D.-H. Kim, N. Sniadecki, Y. Zheng, and all other members of the Gree Research Scholars group for helpful discussion; D. Hailey of the University of Washington (UW) Garvey Imaging Center for his ongoing support and advice; R. Schuck and H. Haeberle for helpful consultation regarding multiphoton microscopy; J. Shadish for providing STEPL vectors for protein expression; and B. Badeau for providing N_3_-*o*NB-RGPQGIWGQGRK(N_3_)-NH_2_. The “Pulmonary arteries (human)” model by finole, the “Seattle Space Needle” model by Intentional3D, and the “Gizmo” model by Multiverse3DDesigns are licensed under Creative Commons licenses and freely available through Thingiverse. We acknowledge support from J. Scott Edgar (In Memoriam) at the UW Mass Spectrometry Center as well as that from the NIH and N. Peters at the UW W. M. Keck Microscopy Center (Grant S10 OD016240). The Thorlabs multiphoton microscope was acquired with and operated under support from the Washington Research Foundation, the UW College of Engineering, the Institute for Stem Cell and Regenerative Medicine, and the Departments of Chemical Engineering, Bioengineering, Chemistry, and Biology. This work was further supported by a generous donation from the Gree Real Estate Foundation; a Faculty Early Career Development (CAREER) Award (DMR 1652141 to C.A.D.), a standard award (DMR 1807398 to C.A.D.), and a Graduate Research Fellowship (DGE 1762114 to R.C.B.) from the National Science Foundation; a seed grant through the NSF-sponsored University of Washington Materials Research Science and Engineering Center (DMR 1719797 to C.A.D.); a Maximizing Investigators’ Research Award (R35GM138036 to C.A.D.) and an additional award (R01DK128551 to K.R.S) from the National Institutes of Health; and an Allen Distinguished Investigator Award, a Paul G. Allen Frontiers Group advised grant of the Paul G. Allen Family Foundation (K.R.S.).

## Availability of Data and Materials

All pertinent experimental procedures, materials, methods, and characterization data are available within this manuscript and its associated Supplementary Information. Plasmids used during the current study are available from the corresponding author on reasonable request.

## References

1. R. Langer, J. P. Vacanti, Tissue Engineering. Science 260, 920–926 (1993).

2. K. Y. Lee, D. J. Mooney, Hydrogels for tissue engineering. Chemical Reviews 101, 1869– 1879 (2001).

3. N. A. Peppas, J. Z. Hilt, A. Khademhosseini, R. Langer, Hydrogels in biology and medicine: From molecular principles to bionanotechnology. Advanced Materials 18, 1345– 1360 (2006).

4. Y. S. Zhang, A. Khademhosseini, Advances in engineering hydrogels. Science 356 (2017).

5. J. Malda, J. Visser, F. P. Melchels, T. Jüngst, W. E. Hennink, W. J. A. Dhert, J. Groll, D. W. Hutmacher, 25th Anniversary Article: Engineering Hydrogels for Biofabrication. Advanced Materials 25, 5011–5028 (2013).

6. S. V. Murphy, A. Atala, 3D bioprinting of tissues and organs. Nat Biotech 32, 773–785 (2014).

7. C. Mandrycky, Z. Wang, K. Kim, D.-H. Kim, 3D bioprinting for engineering complex tissues. Biotechnology Advances 34, 422–434 (2016).

8. R. Levato, T. Jungst, R. G. Scheuring, T. Blunk, J. Groll, J. Malda, From Shape to Function: The Next Step in Bioprinting. Advanced Materials 32, 1906423 (2020).

9. A. Lee, A. R. Hudson, D. J. Shiwarski, J. W. Tashman, T. J. Hinton, S. Yerneni, J. M. Bliley, P. G. Campbell, A. W. Feinberg, 3D bioprinting of collagen to rebuild components of the human heart. Science 365, 482–487 (2019).

10. M. A. Skylar-Scott, J. Mueller, C. W. Visser, J. A. Lewis, Voxelated soft matter via multimaterial multinozzle 3D printing. Nature 575, 330–335 (2019).

11. B. Grigoryan, S. J. Paulsen, D. C. Corbett, D. W. Sazer, C. L. Fortin, A. J. Zaita, P. T. Greenfield, N. J. Calafat, J. P. Gounley, A. H. Ta, F. Johansson, A. Randles, J. E. Rosenkrantz, J. D. Louis-Rosenberg, P. A. Galie, K. R. Stevens, J. S. Miller, Multivascular networks and functional intravascular topologies within biocompatible hydrogels. Science 364, 458–464 (2019).

12. P. N. Bernal, P. Delrot, D. Loterie, Y. Li, J. Malda, C. Moser, R. Levato, Volumetric Bioprinting of Complex Living-Tissue Constructs within Seconds. Advanced Materials 31, 1904209 (2019).

13. U. N. Lee, J. H. Day, A. J. Haack, R. C. Bretherton, W. Lu, C. A. DeForest, A. B. Theberge, E. Berthier, Layer-by-layer fabrication of 3D hydrogel structures using open microfluidics. Lab on a Chip 20, 525–536 (2020).

14. I. S. Kinstlinger, S. H. Saxton, G. A. Calderon, K. V. Ruiz, D. R. Yalacki, P. R. Deme, J. E. Rosenkrantz, J. D. Louis-Rosenberg, F. Johansson, K. D. Janson, D. W. Sazer, S. S. Panchavati, K.-D. Bissig, K. R. Stevens, J. S. Miller, Generation of model tissues with dendritic vascular networks via sacrificial laser-sintered carbohydrate templates. Nature Biomedical Engineering 4, 916–932 (2020).

15. R. Rizzo, D. Ruetsche, H. Liu, M. Zenobi-Wong, Optimized Photoclick (Bio)Resins for Fast Volumetric Bioprinting. Advanced Materials 33, 2102900 (2021).

16. J. A. Brassard, M. Nikolaev, T. Hübscher, M. Hofer, M. P. Lutolf, Recapitulating macro-scale tissue self-organization through organoid bioprinting. Nature Materials 20, 22–29 (2021).

17. E. R. Ruskowitz, C. A. DeForest, Photoresponsive biomaterials for targeted drug delivery and 4D cell culture. Nature Reviews Materials 3, 17087 (2018).

18. T. L. Rapp, C. A. DeForest, Visible Light-Responsive Dynamic Biomaterials: Going Deeper and Triggering More. Advanced Healthcare Materials 9, 1901553 (2020).

19. T. L. Rapp, C. A. DeForest, Targeting drug delivery with light: A highly focused approach. Advanced Drug Delivery Reviews 171 (2021).

20. R. Gharios, R. M. Francis, C. A. DeForest, Chemical and biological engineering strategies to make and modify next-generation hydrogel biomaterials. Matter 6, 4195–4244 (2023).

21. R. M. Francis, C. A. DeForest, 4D Biochemical Photocustomization of Hydrogel Scaffolds for Biomimetic Tissue Engineering. Acc. Mater. Res. 4, 704–715 (2023).

22. B. G. Munoz-Robles, I. Kopyeva, C. A. DeForest, Surface Patterning of Hydrogel Biomaterials to Probe and Direct Cell–Matrix Interactions. Advanced Materials Interfaces 7, 2001198 (2020).

23. M. S. Hahn, L. J. Taite, J. J. Moon, M. C. Rowland, K. A. Ruffino, J. L. West, Photolithographic patterning of polyethylene glycol hydrogels. Biomaterials 27, 2519–2524 (2006).

24. J. M. Karp, Y. Yeo, W. Geng, C. Cannizarro, K. Yan, D. S. Kohane, G. Vunjak-Novakovic, R. S. Langer, M. Radisic, A photolithographic method to create cellular micropatterns. Biomaterials 27, 4755–4764 (2006).

25. S. J. Bryant, K. D. Hauch, B. D. Ratner, Spatial Patterning of Thick Poly(2-hydroxyethyl methacrylate) Hydrogels. Macromolecules 39, 4395–4399 (2006).

26. M. S. Hahn, J. S. Miller, J. L. West, Three-dimensional biochemical and biomechanical patterning of hydrogels for guiding cell behavior. Advanced Materials 18, 2679–2684 (2006).

27. S. H. Lee, J. J. Moon, J. L. West, Three-dimensional micropatterning of bioactive hydrogels via two-photon laser scanning photolithography for guided 3D cell migration. Biomaterials 29, 2962–2968 (2008).

28. C. A. DeForest, B. D. Polizzotti, K. S. Anseth, Sequential click reactions for synthesizing and patterning three-dimensional cell microenvironments. Nature Materials 8, 659–664 (2009).

29. A. M. Kloxin, A. M. Kasko, C. N. Salinas, K. S. Anseth, Photodegradable Hydrogels for Dynamic Tuning of Physical and Chemical Properties. Science 324, 59–63 (2009).

30. R. G. Wylie, S. Ahsan, Y. Aizawa, K. L. Maxwell, C. M. Morshead, M. S. Shoichet, Spatially controlled simultaneous patterning of multiple growth factors in three-dimensional hydrogels. Nature Materials 10, 799–806 (2011).

31. J. C. Culver, J. C. Hoffmann, R. A. Poche, J. H. Slater, J. L. West, M. E. Dickinson, Three-Dimensional Biomimetic Patterning in Hydrogels to Guide Cellular Organization. Advanced Materials 24, 2344–2348 (2012).

32. C. A. DeForest, E. A. Sims, K. S. Anseth, Peptide-functionalized click hydrogels with independently tunable mechanics and chemical functionality for 3D cell culture. Chemistry of Materials 22, 4783–4790 (2010).

33. A. D. Rape, M. Zibinsky, N. Murthy, S. Kumar, A synthetic hydrogel for the high-throughput study of cell–ECM interactions. Nature Communications 6, 8129 (2015).

34. S. C. P. Norris, P. Tseng, A. M. Kasko, Direct Gradient Photolithography of Photodegradable Hydrogels with Patterned Stiffness Control with Submicrometer Resolution. ACS Biomaterials Science & Engineering 2, 1309–1318 (2016).

35. N. Broguiere, I. Lüchtefeld, L. Trachsel, D. Mazunin, R. Rizzo, J. W. Bode, M. P. Lutolf, M. Zenobi-Wong, Morphogenesis Guided by 3D Patterning of Growth Factors in Biological Matrices. Advanced Materials 32, 1908299 (2020).

36. T. A. Pologruto, B. L. Sabatini, K. Svoboda, ScanImage: Flexible software for operating laser scanning microscopes. BioMedical Engineering OnLine 2, 13 (2003).

37. C. A. DeForest, K. S. Anseth, Photoreversible patterning of biomolecules within click-based hydrogels. Angewandte Chemie International Edition 51, 1816–1819 (2012).

38. C. A. DeForest, D. A. Tirrell, A photoreversible protein-patterning approach for guiding stem cell fate in three-dimensional gels. Nature Materials 14, 523–531 (2015).

39. J. C. Grim, T. E. Brown, B. A. Aguado, D. A. Chapnick, A. L. Viert, X. Liu, K. S. Anseth, A Reversible and Repeatable Thiol-Ene Bioconjugation for Dynamic Patterning of Signaling Proteins in Hydrogels. ACS Central Science 4, 909–916 (2018).

40. J. A. Shadish, G. M. Benuska, C. A. DeForest, Bioactive site-specifically modified proteins for 4D patterning of gel biomaterials. Nature Materials 18, 1005–1014 (2019).

41. J. A. Shadish, A. C. Strange, C. A. DeForest, Genetically Encoded Photocleavable Linkers for Patterned Protein Release from Biomaterials. Journal of the American Chemical Society 141, 15619–15625 (2019).

42. R. Warden-Rothman, I. Caturegli, V. Popik, A. Tsourkas, Sortase-tag expressed protein ligation: Combining protein purification and site-specific bioconjugation into a single step. Analytical Chemistry 85, 11090–11097 (2013).

43. J. A. Shadish, C. A. DeForest, Site-Selective Protein Modification: From Functionalized Proteins to Functional Biomaterials. Matter 2, 50–77 (2020).

44. I. Batalov, K. R. Stevens, C. A. DeForest, Photopatterned biomolecule immobilization to guide three-dimensional cell fate in natural protein-based hydrogels. Proceedings of the National Academy of Sciences 118, e2014194118 (2021).

45. H. Krüger, M. Asido, J. Wachtveitl, R. Tampé, R. Wieneke, Sensitizer-enhanced two-photon patterning of biomolecules in photoinstructive hydrogels. Communications Materials 3, 9 (2022).

46. P. E. Farahani, S. M. Adelmund, J. A. Shadish, C. A. DeForest, Photomediated Oxime Ligation as a Bioorthogonal Tool for Spatiotemporally-Controlled Hydrogel Formation and Modification. Journal of Materials Chemistry B 5, 4435–4442 (2017).

47. V. Hagen, B. Dekowski, V. Nache, R. Schmidt, D. Geißler, D. Lorenz, J. Eichhorst, S. Keller, H. Kaneko, K. Benndorf, B. Wiesner, Coumarinylmethyl Esters for Ultrafast Release of High Concentrations of Cyclic Nucleotides upon One-and Two-Photon Photolysis. Angew. Chem. Int. Ed. 44, 7887–7891 (2005).

48. B. C. Campbell, E. M. Nabel, M. H. Murdock, C. Lao-Peregrin, P. Tsoulfas, M. G. Blackmore, F. S. Lee, C. Liston, H. Morishita, G. A. Petsko, mGreenLantern: a bright monomeric fluorescent protein with rapid expression and cell filling properties for neuronal imaging. Proc. Natl. Acad. Sci. U.S.A. 117, 30710–30721 (2020).

49. A. Royant, M. Noirclerc-Savoye, Stabilizing role of glutamic acid 222 in the structure of Enhanced Green Fluorescent Protein. Journal of Structural Biology 174, 385–390 (2011).

50. O. Sarig-Nadir, N. Livnat, R. Zajdman, S. Shoham, D. Seliktar, Laser Photoablation of Guidance Microchannels into Hydrogels Directs Cell Growth in Three Dimensions. Biophysical Journal 96, 4743–4752 (2009).

51. C. A. DeForest, K. S. Anseth, Cytocompatible click-based hydrogels with dynamically tunable properties through orthogonal photoconjugation and photocleavage reactions. Nature Chemistry 3, 925–931 (2011).

52. A. M. Kloxin, K. J. R. Lewis, C. A. DeForest, G. Seedorf, M. W. Tibbitt, V. Balasubramaniam, K. S. Anseth, Responsive culture platform to examine the influence of microenvironmental geometry on cell function in 3D. Integrative Biology 4, 1540–1549 (2012).

53. M. Lunzer, L. Shi, O. G. Andriotis, P. Gruber, M. Markovic, P. J. Thurner, D. Ossipov, R. Liska, A. Ovsianikov, A Modular Approach to Sensitized Two-Photon Patterning of Photodegradable Hydrogels. Angewandte Chemie International Edition 57, 15122–15127 (2018).

54. K. A. Heintz, M. E. Bregenzer, J. L. Mantle, K. H. Lee, J. L. West, J. H. Slater, Fabrication of 3D Biomimetic Microfluidic Networks in Hydrogels. Advanced Healthcare Materials 5, 2153–2160 (2016).

55. N. Brandenberg, M. P. Lutolf, In Situ Patterning of Microfluidic Networks in 3D Cell-Laden Hydrogels. Advanced Materials 28, 7450–7456 (2016).

56. C. K. Arakawa, B. A. Badeau, Y. Zheng, C. A. DeForest, Multicellular Vascularized Engineered Tissues through User-Programmable Biomaterial Photodegradation. Advanced Materials 29, 1703156 (2017).

57. S. G. Rayner, C. C. Howard, C. J. Mandrycky, S. Stamenkovic, J. Himmelfarb, A. Y. Shih, Y. Zheng, Multiphoton-Guided Creation of Complex Organ-Specific Microvasculature. Advanced Healthcare Materials 10, 2100031 (2021).

58. C. Arakawa, C. Gunnarsson, C. Howard, M. Bernabeu, K. Phong, E. Yang, C. A. DeForest, J. D. Smith, Y. Zheng, Biophysical and biomolecular interactions of malaria-infected erythrocytes in engineered human capillaries. Science Advances 6, eaay7243 (2020).

59. T. L. Rapp, C. A. DeForest, Tricolor visible wavelength-selective photodegradable hydrogel biomaterials. Nat Commun 14, 5250 (2023).

60. A. K. Glaser, K. W. Bishop, L. A. Barner, E. A. Susaki, S. I. Kubota, G. Gao, R. B. Serafin, P. Balaram, E. Turschak, P. R. Nicovich, H. Lai, L. A. G. Lucas, Y. Yi, E. K. Nichols, H. Huang, N. P. Reder, J. J. Wilson, R. Sivakumar, E. Shamskhou, C. R. Stoltzfus, X. Wei, A. K. Hempton, M. Pende, P. Murawala, H.-U. Dodt, T. Imaizumi, J. Shendure, B. J. Beliveau, M. Y. Gerner, L. Xin, H. Zhao, L. D. True, R. C. Reid, J. Chandrashekar, H. R. Ueda, K. Svoboda, J. T. C. Liu, A hybrid open-top light-sheet microscope for versatile multi-scale imaging of cleared tissues. Nature Methods 19, 613–619 (2022).

61. O. Chaudhuri, J. Cooper-White, P. A. Janmey, D. J. Mooney, V. B. Shenoy, Effects of extracellular matrix viscoelasticity on cellular behaviour. Nature 584, 535–546 (2020).

