## Supplementary Information for "Grayscale 4D Biomaterial Customization at High Resolution and Scale"

### General synthetic information

Chemical reagents and solvents were purchased from either Sigma-Aldrich or Fisher Scientific and used as received unless otherwise noted. Peptide synthesis reagents were purchased from either ChemPep or Chem-Impex and used as received. Deionized water ($dH_2O$) was generated by a U.S. Filter Corporation Reverse Osmosis System with a Desal membrane. Synthetic chemical reactions were performed under a nitrogen atmosphere in oven-dried glassware and stirred with a Teflon-coated magnetic stir bar unless otherwise noted. Solvents were removed *in vacuo* with a Büchi Rotovapor R-3 equipped with a V-700 vacuum pump and V-855 vacuum controller and a Welch 1400 DuoSeal Belt-Drive high vacuum pump. $^1H$ nuclear magnetic resonance (NMR) data was collected at 298 K on Bruker instruments and chemical shifts are reported relative to tetramethylsilane (TMS, $\delta = 0$). Microwave-assisted peptide synthesis was performed on a CEM Liberty 1. Semi-preparative reversed-phase high-pressure liquid chromatography (RP-HPLC) was performed on a Dionex Ultimate 3000 equipped with a variable multiple wavelength detector, automated fraction collector, and Thermo 5 μm Synchronis silica 250 x 21.2 mm C18 column. Lyophilization was performed on a LABCONCO FreeZone 2.5 Plus freeze-dryer equipped with a LABCONCO rotary vane 117 vacuum pump. Matrix-assisted laser desorption/ionization time of flight (MALDI-TOF) mass spectrometry was performed in reflectron positive ion mode or reflectron negative ion mode on a Bruker AutoFlex II using a matrix of α-cyano-4-hydroxycinnamic acid:2,5-dihydroxy benzoic acid (2:1). Whole-protein mass spectrometry was performed using a Waters Synapt – G2 QTOF. The light source for the photochemical cleavage was a Lumen Dynamics OmniCure S1500 Spot UV Curing system with an internal 365 nm filter and an external 360 nm cut-on long-pass filter. Light intensity was measured using a Cole-Parmer Radiometer (Series 9811-50, $\lambda = 365$ nm). Confocal microscopy was performed at the University of Washington Keck Microscopy Center on a Leica SP8X confocal microscope or in the DeForest lab on a Leica Stellaris 5 microscope. Multiphoton lithography was performed on an Olympus FV1000 MPE BX61 Multiphoton Microscope at the Garvey Imaging Center at the University of Washington or on a Thorlabs Bergamo II multiphoton microscope in the DeForest lab. Polymerase chain reaction (PCR) was performed in a Bioer LifeECO thermal cycler. Protein expression was performed in a Thermo Scientific MaxQ 4000 shaker incubator. Cells were lysed using a Fisher Scientific Model 505 Sonic Dismembrator with a 1.27 cm diameter probe. Drone footage was acquired using a DJI Mavic Mini.

### Synthesis of previously reported compounds used in this work

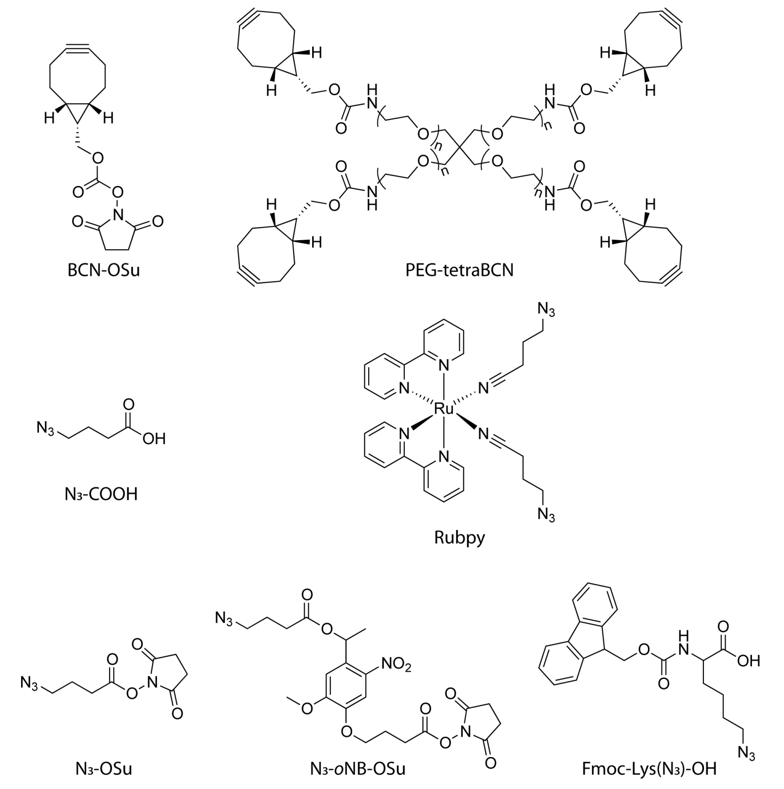

(1R,8S,9s)-bicyclo[6.1.0]non-4-yn-9-ylmethyl (2,5-dioxopyrrolidin-1-yl) carbonate (BCN-OSu), poly(ethylene glycol) tetrabicyclononyne (PEG-tetraBCN, $M_n \sim 20{,}000$ Da), 4-azidobutanoic acid ($N_3$-COOH), 2,5-dioxopyrrolidin-1-yl 4-azidobutanoate ($N_3$-OSu), 2,5-dioxopyrrolidin-1-yl 4-(4-(1-((4-azidobutanoyl)oxy)ethyl)-2-methoxy-5-nitrophenoxy)butanoate ($N_3$-*o*NB-OSu), $N^2$-(((9*H*-fluoren-9-yl)methoxy)carbonyl)-$N^6$-diazolysine (Fmoc-Lys($N_3$)-OH), and Ru(2,2'-bipyridine)$_2$(4-azidobutanenitrile)$_2$ (Rubpy) were synthesized as previously reported[1–3].

**Method S1 GIZMO photopatterning**

GIZMO was implemented on both an Olympus multiphoton microscope (Olympus FV1000 MPE BX61) powered by FluoView software and a Thorlabs system (Bergamo II) driven by ScanImage®.

To facilitate widespread adoption of these techniques and in partnership with Vidrio Technologies, GIZMO is now included within the current distribution of ScanImage® as the "Print3D" module.

Code to implement GIZMO using FluoView is available at: https://github.com/deforestgroup.

Data in main text Figures 2C-E, 3E-F, and 3H, as well as Supplementary Figure S3, were generated using FluoView on the Olympus microscope; all other patterning experiments were performed using ScanImage® on the Thorlabs system.

Specific patterning conditions are given throughout the manuscript body and supporting information.

**Method S2 Synthesis of PEG-diazide ($N_3$-PEG-$N_3$)**

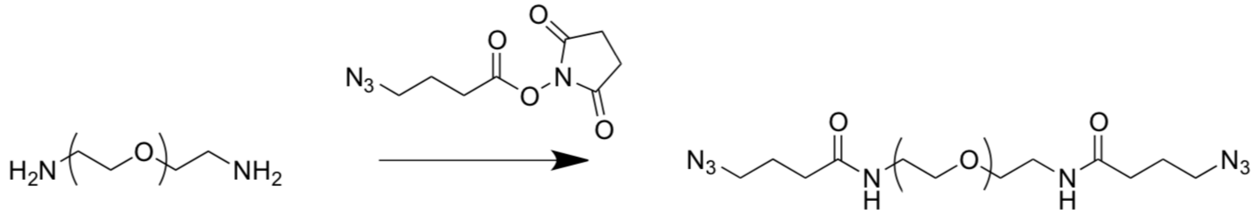

Linear poly(ethylene glycol) diamine ($M_n$ ~ 3,500 Da, 1 g, 0.57 mmol $NH_2$, 1x, Jenkem) and $N_3$-OSu (194 mg, 0.86 mmol, 1.5x) were dissolved in dimethylformamide (5 mL). *N*,*N*-Diisopropylethylamine (398 µL, 294 mg, 2.28 mmol, 4x) was added to the mixture, and the reaction was stirred overnight, diluted in water (15 mL), dialyzed (MWCO ~ 2 kDa, SpectraPor), and lyophilized to yield a white powder (1.00 g, quantitative yield). $^1$H NMR (500 MHz, $CDCl_3$) δ 3.75 (m, 4H), 3.65-3.61 (m, 318H), 3.28 (m, 4H), 2.35 (m, 4H), 1.86 (m, 4H). Functionalization was found to be >95% by comparing integral values for hydrogens introduced upon azide coupling (δ 3.28, 2.35, 1.86) with those from the PEG backbone (δ 3.60-3.42).

### Method S3 Plasmid construction for protein expression and STEPL purification

STEPL plasmids for used for expression of C-terminally modified mCherry, EGFP and mGreenLantern (mGL), as previously reported[4,5]. The DNA open reading frame sequences (5' → 3') for these systems are shown below. Nucleotides shown in black correspond to that for the specific protein of interest. Bases shown in orange correspond to STEPL portion common to all constructs, which features the C-terminal LPETG sortase recognition motif, a flexible $(GGS)_5$ linker, SrtA, and a 6xHis tag.

*mCherry-STEPL*:

ATGGTTTCCAAGGGCGAAGAAGACAACATGGCGATCATCAAAGAATTTATGCGTTTTAAAGTTCACATGGAAGGTTC
TGTTAACGGTCATGAGTTCGAAATTGAAGGTGAGGGTGAAGGTCGCCCGTACGAAGGTACCCAGACCGCGAAACTGA
AAGTTACCAAAGGTGGTCCGCTGCCGTTCGCGTGGGACATCCTCAGCCCGCAGTTCATGTACGGTTCTAAAGCGTAC
GTTAAACATCCGGCGGACATTCCAGACTACCTCAAACTCTCTTTCCCTGAAGGTTTCAAATGGGAACGTGTTATGAA
CTTCGAGGACGGTGGTGTTGTCACCGTTACCCAGGACTCTTCTCTGCAGGACGGCGAGTTCATCTACAAGGTCAAAC
TGCGTGGCACCAACTTCCCGTCTGACGGTCCGGTTATGCAGAAAAAAACCATGGGTTGGGAAGCGTCTTCTGAACGT
ATGTACCCGGAAGATGGTGCGCTGAAAGGCGAAATCAAACAGCGTCTGAAGCTCAAAGACGGCGGTCACTACGACGC
GGAGGTTAAAACCACCTACAAAGCGAAAAAGCCGGTTCAACTGCCGGGTGCGTACAACGTTAATATCAAGCTGGACA
TCACCTCTCACAACGAAGACTACACCATCGTTGAACAGTACGAACGTGCGGAAGGCCGTCACTCTACCGGTGGTATG
GACGAACTGTACAAGCTCGAGCTGCCGGAAACCGGTGGTGGTAGTGGTGGCTCTGGCGGTTCTGGTGGCAGTGGCGG
TAGCCAAGCTAAACCTCAAATTCCGAAAGATAAATCAAAAGTGGCAGGCTATATTGAAATTCCAGATGCTGATATTA
AAGAACCAGTATATCCAGGACCAGCAACACCTGAACAATTAAATAGAGGTGTAAGCTTTGCAGAAGAAAATGAATCA
CTAGATGATCAAAATATTTCAATTGCAGGACACACTTTCATTGACCGTCCGAACTATCAATTTACAAATCTTAAAGC
AGCCAAAAAAGGTAGTATGGTGTACTTTAAAGTTGGTAATGAAACACGTAAGTATAAAATGACAAGTATAAGAGATG
TTAAGCCAACAGATGTAGAAGTTCTAGATGAACAAAAAGGTAAAGATAAACAATTAACATTAATTACTTGTGATGAT
TACAATGAAAAGACAGGCGTTTGGGAAAAACGTAAAATCTTTGTAGCTACAGAAGTCAAACATCACCACCATCATCA
CTAA

*EGFP-STEPL*:

ATGGTGAGCAAGGGCGAGGAGCTGTTCACCGGGGTGGTGCCCATCCTGGTCGAGCTGGACGGCGACGTAAACGGCCA
CAAGTTCAGCGTGTCCGGCGAGGGCGAGGGCGATGCCACCTACGGCAAGCTGACCCTGAAGTTCATCTGCACCACCG
GCAAGCTGCCCGTGCCCTGGCCCACCCTCGTGACCACCCTGACCTACGGCGTGCAGTGCTTCAGCCGCTACCCCGAC
CACATGAAGCAGCACGACTTCTTCAAGTCCGCCATGCCCGAAGGCTACGTCCAGGAGCGCACCATCTTCTTCAAGGA
CGACGGCAACTACAAGACCCGCGCCGAGGTGAAGTTCGAGGGCGACACCCTGGTGAACCGCATCGAGCTGAAGGGCA
TCGACTTCAAGGAGGACGGCAACATCCTGGGGCACAAGCTGGAGTACAACTACAACAGCCACAACGTCTATATCATG
GCCGACAAGCAGAAGAACGGCATCAAGGTGAACTTCAAGATCCGCCACAACATCGAGGACGGCAGCGTGCAGCTCGC
CGACCACTACCAGCAGAACACCCCCATCGGCGACGGCCCCGTGCTGCTGCCCGACAACCACTACCTGAGCACCCAGT
CCGCCCTGAGCAAAGACCCCAACGAGAAGCGCGATCACATGGTCCTGCTGGAGTTCGTGACCGCCGCCGGGATCACT
CTCGGCATGGACGAGCTGTACAAGCTCGAGCTGCCGGAAACCGGTGGTGGTAGTGGTGGCTCTGGCGGTTCTGGTGG
CAGTGGCGGTAGCCAAGCTAAACCTCAAATTCCGAAAGATAAATCAAAAGTGGCAGGCTATATTGAAATTCCAGATG
CTGATATTAAAGAACCAGTATATCCAGGACCAGCAACACCTGAACAATTAAATAGAGGTGTAAGCTTTGCAGAAGAA
AATGAATCACTAGATGATCAAAATATTTCAATTGCAGGACACACTTTCATTGACCGTCCGAACTATCAATTTACAAA
TCTTAAAGCAGCCAAAAAAGGTAGTATGGTGTACTTTAAAGTTGGTAATGAAACACGTAAGTATAAAATGACAAGTA
TAAGAGATGTTAAGCCAACAGATGTAGAAGTTCTAGATGAACAAAAAGGTAAAGATAAACAATTAACATTAATTACT
TGTGATGATTACAATGAAAAGACAGGCGTTTGGGAAAAACGTAAAATCTTTGTAGCTACAGAAGTCAAACATCACCA
CCATCATCACTAA

*mGL-STEPL*:

ATGGTCAGCAAGGGCGAAGAATTGTTCACAGGCGTCGTCCCGATCTTAGTTGAGCTGGATGGTGACGTTAACGGCCA
TAAATTTAGCGTACGGGGAGAGGGTGAGGGTGACGCTACAAATGGCAAGTTGACACTCAAATTCATCTGTACCACCG
GGAAGTTGCCGGTTCCGTGGCCTACGCTCGTCACCACGTTAGGGTATGGAGTAGCATGTTTCGCAAGATACCCGGAC
CACATGAAGCAGCACGATTTTTTCAAAAGTGCCATGCCCGAGGGTTACGTACAGGAGCGTACGATATCCTTCAAAGA
TGACGGGACCTATAAGACTCGGGCTGAGGTCAAGTTCGAAGGCGACACGTTAGTTAACCGCATTGTTTTAAAGGGGA
TAGACTTTAAGGAAGACGGCAACATTCTCGGGCATAAACTGGAGTACAACTTCAATAGCCACAAGGTCTACATCACC

GCGGACAAACAGAAGAACGGCATAAAAGCCAACTTCAAGACCCGGCATAATGTGGAGGACGGAGGCGTGCAGTTGGC
AGATCACTATCAACAAAATACACCTATTGGTGACGGCCCCGTTTTGTTGCCGGATAACCACTACCTGTCCCACCAAA
GTAAGTTGAGCAAAGATCCCAACGAGAAACGTGACCATATGGTTTTAAAGGAGAGAGTGACCGCGGCCGGCATCACA
CACGACATGGATGAGTTGTACAAACTCGAGCTGCCGGAAACCGGTGGTGGTAGTGGTGGCTCTGGCGGTTCTGGTGG
CAGTGGCGGTAGCCAAGCTAAACCTCAAATTCCGAAAGATAAATCAAAAGTGGCAGGCTATATTGAAATTCCAGATG
CTGATATTAAAGAACCAGTATATCCAGGACCAGCAACACCTGAACAATTAAATAGAGGTGTAAGCTTTGCAGAAGAA
AATGAATCACTAGATGATCAAAATATTTCAATTGCAGGACACACTTTCATTGACCGTCCGAACTATCAATTTACAAA
TCTTAAAGCAGCCAAAAAAGGTAGTATGGTGTACTTTAAAGTTGGTAATGAAACACGTAAGTATAAAATGACAAGTA
TAAGAGATGTTAAGCCAACAGATGTAGAAGTTCTAGATGAACAAAAAGGTAAAGATAAACAATTAACATTAATTACT
TGTGATGATTACAATGAAAAGACAGGCGTTTGGGAAAAACGTAAAATCTTTGTAGCTACAGAAGTCAAACACCATCA
TCATCACCATTAA

**Method S4 Fmoc solid-phase peptide synthesis**

A CEM Liberty1 was used to perform microwave-assisted Fmoc solid-phase peptide synthesis (SPPS, 0.25 mmol scale). Fmoc deprotection was performed in 20% piperidine (v/v) in dimethylformamide (DMF) with 1-hydroxybenzotriazole (HOBt, 0.1 M, 90 °C, 90 sec). Amino acids were coupled to resin-bound peptides upon treatment (75 °C, 5 min) with Fmoc-protected amino acid (2 mmol, 4x), 2-(1H-benzotriazol-1-yl)-1,1,3,3-tetramethyluronium hexafluorophosphate (HBTU, 2 mmol, 4x), and *N*,*N*-diisopropylethylamine (DIEA, 2 mmol, 4x) in a mixture of DMF (9 mL) and *N*-Methyl-2-pyrrolidone (NMP, 2 mL). Room-temperature couplings were performed without microwave assistance (1 hr).

**Method S5 Synthesis of H-GGGGDDK(*o*NB-$N_3$)-$NH_2$**

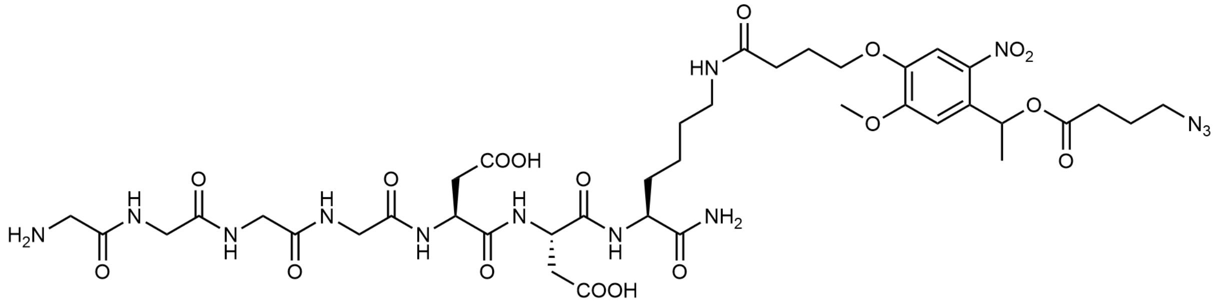

The resin-bound peptide Boc-GGGGDDK(Mtt)-$NH_2$ was synthesized as previously reported[4] by Fmoc SPPS (Method S2) on Rink amide resin (0.25 mmol scale). The resin was washed with DMF (3x) and dichloromethane (DCM, 3x) prior to Mtt cleavage (2x, 9 min, 15 mL, 97:2:1 DCM:TIS:TFA). Resin was washed (DCM, 3x; DMF, 3x) prior to treatment (2 hr) with $N_3$-*o*NB-OSu (1.5x, 0.375 mmol, 90 mg) and DIEA (4x, 1 mmol, 87 µL) in minimal DMF. Resin was washed (DMF, 3x; DCM, 3x) prior to peptide cleavage/deprotection (95:5 TFA:$H_2O$, 20 mL, 2 hr) and precipitation (diethyl ether, 180 mL, 0 °C, 2x). The crude peptide was purified *via* RP-HPLC using a 55-minute gradient from 5-100% acetonitrile:$H_2O$; lyophilization yielded the final product (H-GGGGDDK(*o*NB-$N_3$)-$NH_2$) as a yellow solid (70 mg, 0.070 mmol, 27% overall yield). Peptide purity was confirmed using MALDI-TOF: calculated for $C_{39}H_{56}N_{13}O_{18}^-$ [M - $^1$H]$^-$, 994.39; observed 993.47.

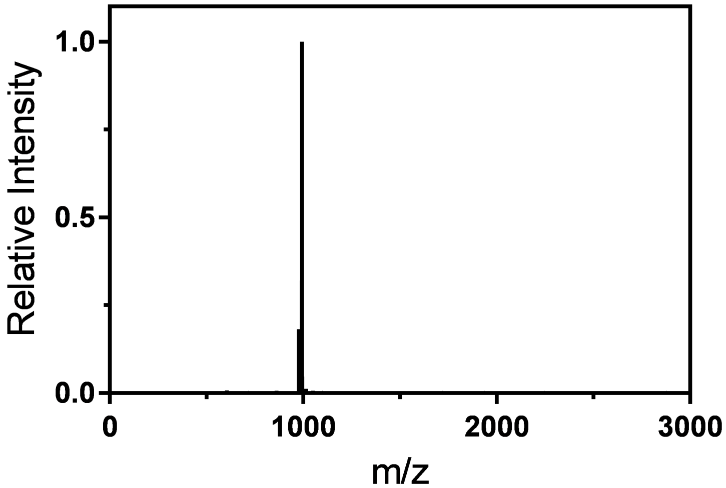

**Method S6 Synthesis of H-GGGGDDK(CHO)-$NH_2$**

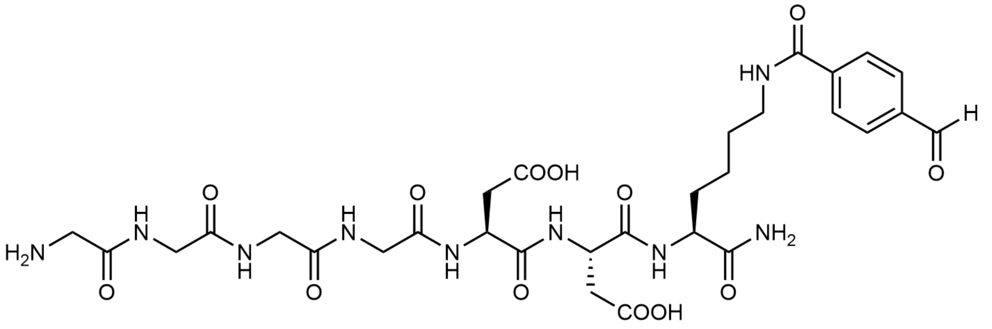

The resin-bound peptide Boc-GGGGDDK(Mtt)-$NH_2$ was synthesized as previously reported[4] by Fmoc SPPS (Method S2) on Rink amide resin (0.25 mmol scale). The resin was washed with DMF (3x) and dichloromethane (DCM, 3x) prior to Mtt cleavage (2 min, 15 mL, 97:2:1 DCM:TIS:TFA, 9x). Resin was washed (DCM, 3x; DMF, 3x) prior to treatment (1 hr) with 4-formylbenzoic acid (4x, 1 mmol, 150 mg) by HATU coupling (3.95x, 0.988 mmol, 188 mg) and DIEA (8x, 2 mmol, 174 µL) in minimal DMF. Resin was washed (DMF, 3x; DCM, 3x) prior to peptide cleavage/deprotection (95:5 TFA:$H_2O$, 20 mL, 2 hr) and precipitation (diethyl ether, 180 mL, 0 °C, 2x). The crude peptide was purified *via* RP-HPLC using a 55-minute gradient from 5-100% acetonitrile:$H_2O$; lyophilization yielded the final product (H-GGGGDDK(CHO)-$NH_2$) as a white solid (39 mg, 0.053 mmol, 21% overall yield). Peptide purity was confirmed using MALDI-TOF: calculated for $C_{30}H_{42}N_9O_{13}^+$ $[M + {}^1H]^+$, 736.29; observed 736.39.

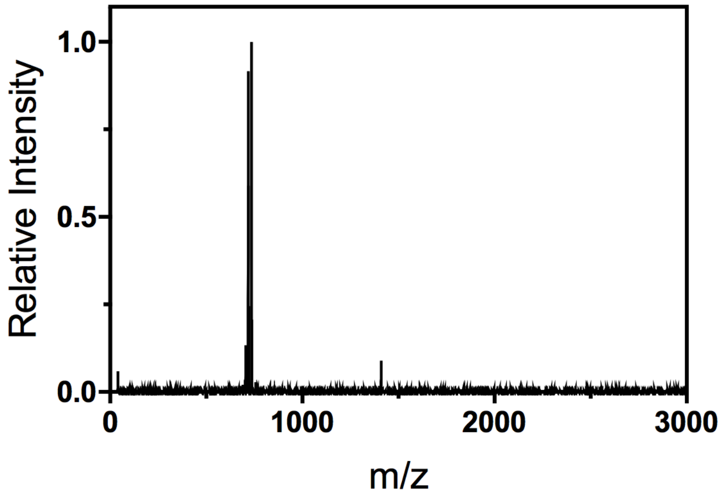

### Method S7 Protein expression and purification by STEPL

STEPL plasmids for mCherry, EGFP, and mGL were transformed into BL21(DE3) *Escherichia coli* (Thermo Fisher). Transformants were grown at 37 °C in lysogeny broth containing ampicillin (100 µg $mL^{-1}$) until reaching an optical density of 0.6 (λ = 600 nm). Isopropyl ß-D-1-thiogalactopyranoside was added (0.5 mM final concentration) to induce culture, prior to overnight expression at 18 °C.

Cells were harvested *via* centrifugation (7,000 g, 10 min). The cell pellet was resuspended in lysis buffer (40 mL, 20 mM Tris, 50 mM NaCl, 10 mM imidazole, 1 mM phenylmethylsulfonyl fluoride) and sonicated on ice (6 cycles of 3 minutes at 30% amplitude 33% duty cycle and 3 min resting). Soluble and insoluble fractions were separated *via* centrifugation (5,000 g, 20 min).

Clarified lysate was applied to Ni-NTA resin (2.5 mL) and incubated under mild agitation (4 °C, 1 hr). The flow-through was discarded, and the resin was washed with wash buffer (20 mM Tris, 50 mM NaCl, 20 mM imidazole, 20 mL, 5x) and STEPL buffer (20 mM Tris, 50 mM NaCl, 20 mL, 1x). The polyglycine probe (20 molar excess) was added to resin in conjugation buffer (20 mM Tris, 50 mM NaCl, 100 µM $CaCl_2$, 2 mL) to promote intramolecular sortagging (37 °C, 4 hr).

The conjugated protein solution was collected, and the resin was washed with STEPL buffer (1 mL, 5x) to collect any remaining protein. The protein solution was dialyzed against STEPL buffer using ThermoFisher SnakeSkin Dialysis Tubing (molecular weight cut-off, MWCO ~ 10 kDa) to remove any unconjugated peptide and concentrated using an Amicon centrifugal spin column (MWCO ~ 10 kDa). Typical yields for purified proteins following STEPL were ~15 mg per liter of cell culture.

mCherry was sortagged with H-GGGGDDK(*o*NB-$N_3$)-$NH_2$ (yielding mCherry-*o*NB-$N_3$). EGFP and mGL were sortagged with H-GGGGDDK(CHO)-$NH_2$ (respectively yielding EGFP-CHO and mGL-CHO).

Sortagged protein purity was verified by LC/MS, with observed molecular masses matching those expected for all species:

> mCherry-*o*NB-$N_3$: expected 28,285 and 28,154 Da (-Met); observed 28,284 Da and 28,154 Da (-Met)
> EGFP-CHO: 28,324 Da; observed 28,321 Da
> mGL-CHO: expected 28,205 Da; observed 28,205 Da

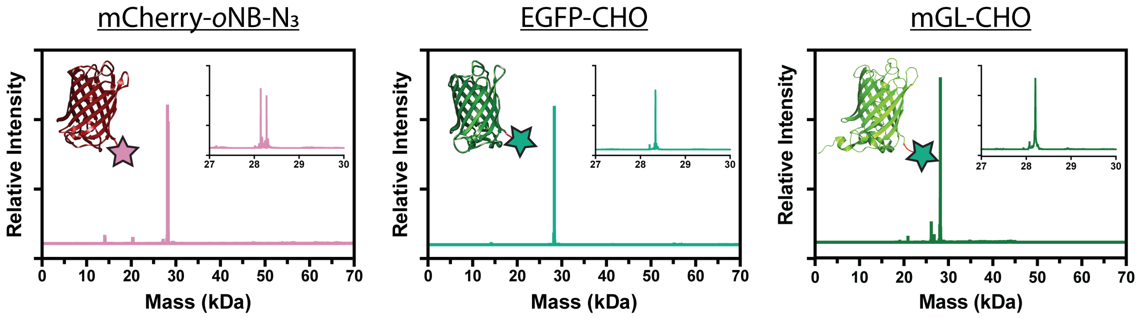

**Method S8 Hydrogel synthesis for biomolecule photorelease studies**

PEG-tetraBCN ($M_n \approx$ 20 kDa, 2 mM) was combined with $N_3$-PEG-$N_3$ ($M_n \approx$ 3.5 kDa, 4 mM) and either mCherry-*o*NB-$N_3$ (100 μM) or $N_3$-*o*NB-GRGDSK(AF488)-$NH_2$ (100 μM) in PBS. Gelation was allowed to proceed for 1 h between Rain-X-treated glass slides with silicone rubber spacers (0.5 mm thick, McMaster-Carr). Following slide separation, gels were equilibrated overnight in PBS prior to use.

**Figure S1 Patterning time scales with complexity in prior approaches, but not in GIZMO**

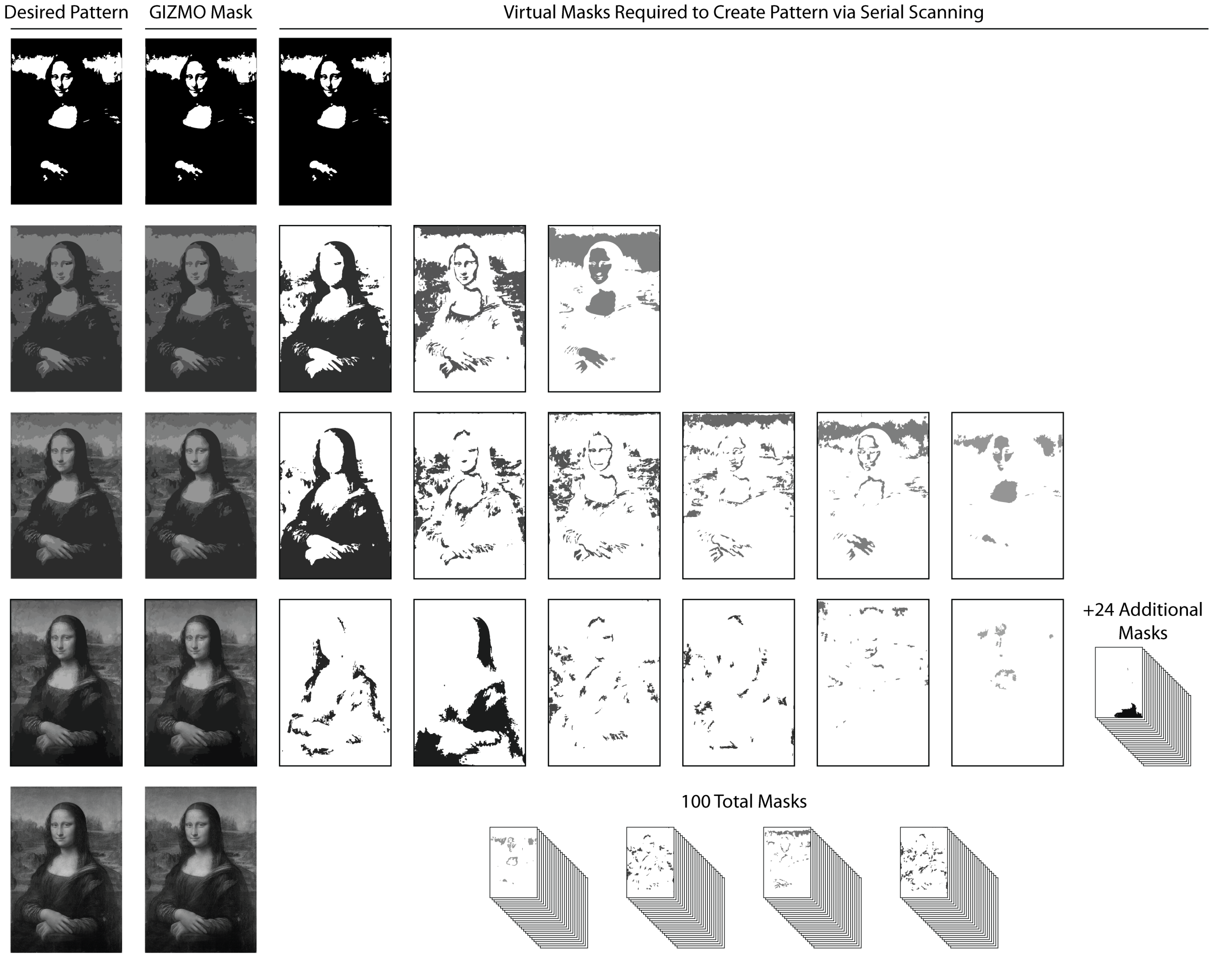

Grayscale patterning afforded recently relies on serial scanning of virtual "masks", each comprised of an assemblage of regions of interests with a shared pixel intensity, at the same gel z-location with distinct laser powers[6]. Though highly detailed patterning may be possible, the number of masks used at a given z-position (and therefore patterning times) scales directly with image intensity depth. Since the GIZMO method specifies laser power throughout continuous raster scanning on a voxel-by-voxel basis, only a single mask is required at each z-position regardless of image complexity, shortening overall patterning times by several orders of magnitude.

### Figure S2 Assessing pattern fidelity via parity analysis

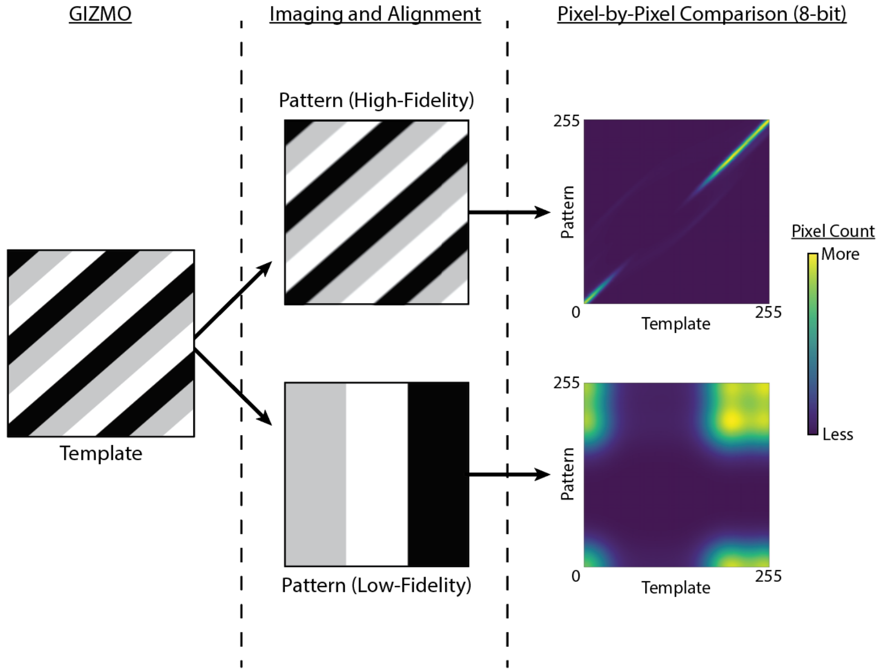

To assess pattern fidelity, GIZMO input template files and confocal images from patterned gels were manually cropped and aligned in Adobe Illustrator. Using ImageJ, the confocal image resolution and min/max pixel values were normalized to the template. Adjusted files were then analyzed in Python by plotting the intensities of each pixel pair as a heatmap with identical values falling on the y = x diagonal. Pixel-by-pixel fidelity is visualized as a pairwise comparison of intensities between equivalently located pixels where a 'perfect' reproduction of the template is plotted along y = x.

Large images were binned by pixel intensity, then each bin was equally and randomly sampled to prevent underrepresented intensity regions from being buried in the heatmap. In cases where only a few pixels fell within a given bin, the entire bin was sampled and any resulting gap in the heatmap deemed acceptable.

**Method S9 Synthesis of $N_3$-*o*NB-GRGDSK(AF488)-$NH_2$**

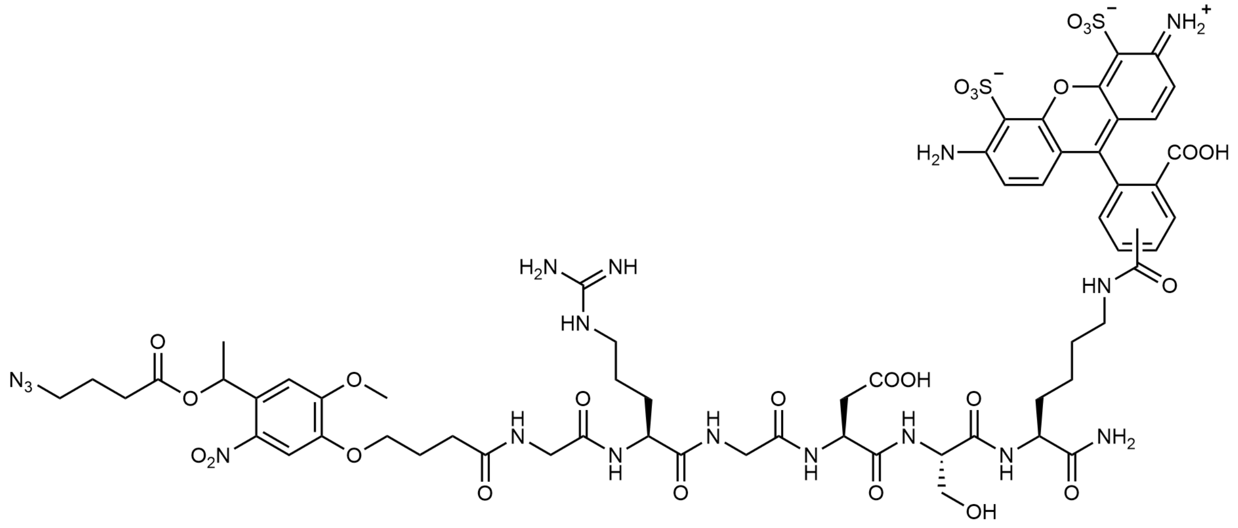

The peptide Fmoc-GRGDSK(boc)-$NH_2$ was synthesized by SPPS. Following N-terminal Fmoc deprotection, the resin was rinsed thrice with DMF, then coupled to $N_3$-*o*NB-OH (205 mg, 0.5 mmol, 2x molar excess) for 90 minutes at room temperature using HATU (188 mg, 0.495 mmol, 1.975x molar excess) in minimal (4 mL) DMF with DIEA (129 mg, 174 µL, 1 mmol, 4x molar excess) as a base. Conjugation of the photolabile azide was verified by the kaiser test. The peptide was then treated with cleavage cocktail (95% TFA 2.5% TIS, 2.5% water) for 180 minutes, precipitated in ice cold diethyl ether, and purified by reverse-phase HPLC on a 45-minute gradient from 20% – 100% acetonitrile in water containing TFA (0.1%), yielding 8.4 mg peptide (1009.8 Da expected, 1010.345 Da observed by MALDI mass spectrometry, secondary peak corresponds to photocleavage product accompanying MALDI ionization, 3.2% overall yield).

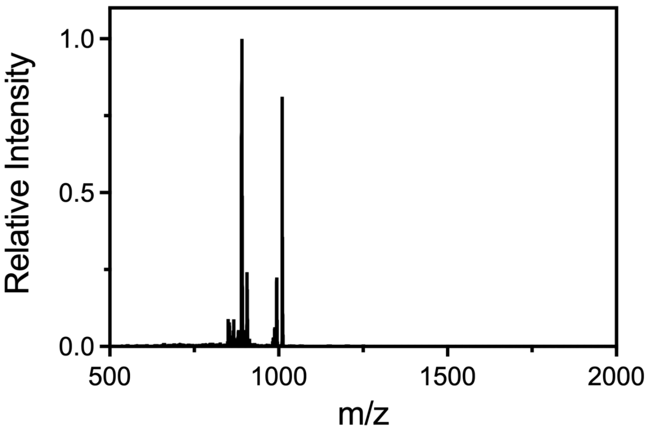

Peptide was then resuspended in DMSO at 50 mM concentration and labeled with Alexa Fluor 488 NHS Ester (Thermo Fisher, 10 mM in DMSO) at a ratio of 1:50 (10 µL peptide, 1 µL AF488-NHS, 9 µL PBS) overnight at room temperature.

The fluorescently labeled peptide [denoted $N_3$-*o*NB-GRGDSK(AF488)-$NH_2$] was used without additional purification.

### Method S10 Synthesis of DCMAC-HNO-TEG-BCN

*Synthesis of 2,2'-[({oxybis[ethane-2,1-diyl]}bis[oxy])bis(ethane-2,1-diyl)]bis(oxy)bis(isoindoline-1,3-dione) **2***

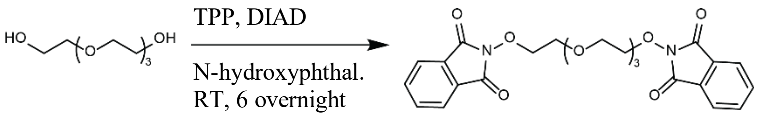

**2**

**2** was synthesized as previously reported with minor changes[7]. Tetraethylene glycol (5.0 g, 26 mmol), triphenylphosphine (14.2 g, 54 mmol), *N*-hydroxyphthalimide (9.3 g, 57 mmol) were added to a flame-dried round bottom flask before purging with $N_2$ gas for 5 minutes. Reactants were suspended in 100 mL of dry dichloromethane, then diisopropyl azodicarboxylate (10.9 g, 54 mmol) was added slowly, allowing the dark orange color to fade before each addition. The reaction was left to stir overnight at room temperature, then concentrated *in vacuo* and diluted with 500 mL diethyl ether. After stirring vigorously for several hours on ice, the white precipitate was collected by filtration, then recrystallized from diethyl ether in minimal dichloromethane. At this point, a small amount of red byproduct also crystallized which was removed by redissolving solids in dichloromethane and washing twice each with water and brine before drying over sodium sulfate. The organics were concentrated *in vacuo* and recrystallized again from diethyl ether to yield 5.8 g of **2** (12 mmol, 47%), a colorless solid which was used without further purification. $^1$H NMR (500 MHz, $CDCl_3$): δ = 3.47-3.53 (m, 4 H), 3.58-3.65 (m, 4 H), 3.80-3.89 (m, 4 H), 4.31-4.41 (m, 4 H), 7.72-7.78 (m, 4 H), 7.80-7.86 (m, 4 H). MS (ESI-MS): calculated for $C_{24}H_{28}N_3O_9$ ($[M+NH_4]^+$): 502.182; found 502.183.

*Synthesis of 2,2'-[({oxybis[ethane-2,1-diyl]}bis[oxy])bis(ethane-2,1-diyl)]bis(hydroxylamine) (TEG-diOA) **3***

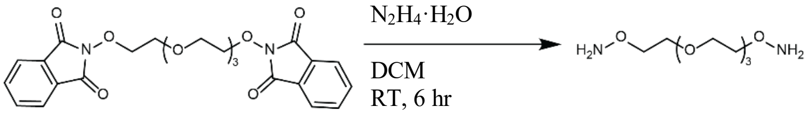

**2** **3**

**3** was synthesized as previously reported[7]. A solution of **2** (5.8 g, 12.0 mmol) was prepared in 100 mL of dichloromethane. Hydrazine monohydrate (1.3 g, 26 mmol, 2.1 eq.) was added, and the reaction was stirred vigorously for 6 hours at room temperature. The off-white precipitate was removed by filtration, then the solution was concentrated *in vacuo* and diluted with diethyl ether. The solution was stirred on ice for 15 minutes, then filtered and concentrated *in vacuo*. The residue was dissolved in chloroform and stirred on ice for 5 minutes, then filtered, concentrated, and filtered again. A final solvent evaporation yielded 2.3 g (11.8 mmol, 85%) of **3** as a clear and slightly yellow oil. $^1$H NMR (500 MHz, $CDCl_3$): δ = 3.63-3.71 (m, 12 H), 3.82-3.86 (m, 4 H), 5.44 (br. s, 4 H). MS (ESI-MS): calculated for $C_8H_{21}N_2O_5$ ($[M+H]^+$): 225.145; found 225.145.

*Synthesis of 7-[bis(tert-butoxycarbonylmethyl)amino]-4-methylcoumarin (DCMAC) **4***

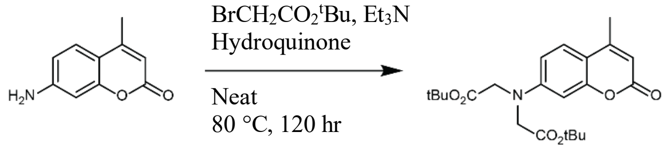

**4**

**4** was synthesized as previously reported.[8] 7-amino-4-methylcoumarin (15 g, 85.6 mmol), *tert*-butyl bromoacetate (45 mL, ), hydroquinone (1.5 g, mmol), and diisopropylethylamine (45 mL, 258.3 mmol) were combined in a round bottom flask and stirred at 80 °C for 5 days in the dark. The slushy product was dissolved in dichloromethane and washed once with water and once with brine, then dried over $MgSO_4$. Solvent was removed *in vacuo*, and the crude product was purified by silica gel chromatography using 1:2 ethyl acetate:hexanes ($R_f$ = 0.6, 1:1 ethyl acetate:hexanes) to yield 22.3 g (55.3 mmol, 65%) of **4**, a white crystalline solid. $^1H$ NMR (500 MHz, $CDCl_3$): δ = 1.47 (s, 18 H), 2.32-2.37 (d, J = 0.8 Hz, 3 H), 4.05 (s, 4 H), 6.00-6.03 (d, J = 1.0 Hz, 1 H), 6.43-6.46 (d, J = 2.5 Hz, 1 H), 6.50-6.54 (dd, J = 2.7 Hz, 8.9 Hz), 7.40-7.44 (d, J = 8.9 Hz, 1 H). MS (ESI-MS): calculated for $C_{22}H_{30}NO_6$ ($[M+H]^+$): 404.207; found: 404.208.

*Synthesis of 7-[bis(tert-butoxycarbonylmethyl)amino]-4-(hydroxymethyl)coumarin* ***5***

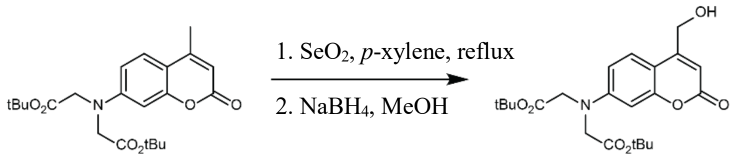

**4** **5**

**5** was synthesized as previously reported.[8] **4** (22.3 g, 55.3 mmol) and selenium dioxide (12.3 g, 110.9 mmol) were dissolved in 250 mL of dry *p*-xylene in a flame-dried round bottom flask. The solution was refluxed for 5.5 hours, then hot filtered to remove tars. Solvent was removed *in vacuo*, and the resulting yellow solids were suspended in 400 mL of methanol. Sodium borohydride (4.36 g, 115.3 mmol) was added slowly, then allowed to react for an additional 30 minutes at room temperature. The pH was adjusted to 2 using 1 M HCl, then diluted with water and extracted 4x with dichloromethane. The combined organics were washed with brine and dried over anhydrous sodium sulfate. Solvent was removed *in vacuo*, and the crude product was purified by silica gel chromatography using 1:1 ethyl acetate:hexanes ($R_f$ = 0.2, 1:1 ethyl acetate:hexanes) to yield 9.7 g of **5** (23.2 mmol, 42%), a yellow crystalline solid. $^1H$ NMR (500 MHz, $CDCl_3$): δ = 1.48 (s, 18 H), 2.02-2.08 (t, J = 6.1 Hz, 1 H), 4.05 (s, 4 H), 4.78-4.84 (dd, J = 1.4 Hz, 5.9 Hz), 6.32-6.35 (t, J = 1.4 Hz, 1 H), 6.45-6.47 (d, J = 2.6 Hz, 1 H), 6.47-6.51 (dd, J = 2.8 Hz, 8.9 Hz), 7.30-7.34 (d, J = 8.8 Hz, 1 H). MS (ESI-MS): calculated for $C_{22}H_{30}NO_7$ ($[M+H]^+$): 420.202; found: 404.200.

*Synthesis of 7-[bis(tert-butoxycarbonylmethyl)amino]-4-coumarinylmethyl (2,5-dioxo-1-pyrrolidinyl) carbonate **6***

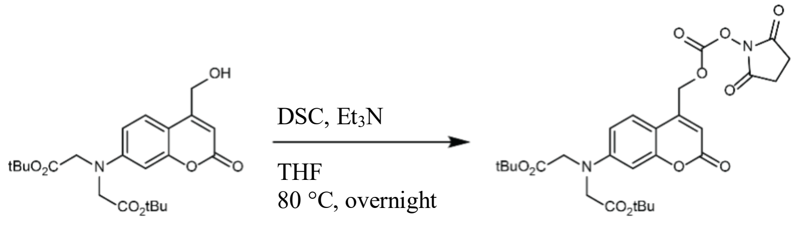

**5** **6**

**5** (4.0 g, 9.5 mmol) and disuccinimidyl carbonate (5 g, 19.5 mmol) were dissolved in 80 mL of dry THF in a flame-dried round bottom flask purged with $N_2$. Triethylamine (3.63 g, 35.9 mmol) was added, and the reaction was allowed to continue overnight at room temperature in the dark. The reaction was diluted with 1:1 ethyl acetate:diethyl ether, then washed 3x with water and once with brine before being dried over anhydrous magnesium sulfate and filtered. Solvent was removed *in vacuo*, and the dark yellow crude product was purified by silica gel chromatography using 1:1 ethyl acetate:hexanes ($R_f$ = 0.12, 1:1 ethyl acetate:hexanes) to yield 3.97 g of **6** (7.1 mmol, 75%), a yellow solid. $^1$H NMR (500 MHz, $CDCl_3$): δ = 1.48 (s, 18 H), 2.87 (s, 4 H), 4.08 (s, 4 H), 5.43 (s, 2 H), 6.33 (s, 1 H), 6.48-6.50 (d, J = 2.4 Hz, 1 H), 6.54-6.59 (dd, J = 2.5 Hz, 9.0 Hz), 6.74-6.93 (br. s, 1 H), 7.32-7.35 (d, J = 8.9 Hz, 1 H). MS (ESI-MS): calculated for $C_{27}H_{33}N_2O_{11}$ ($[M+H]^+$): 561.208; found 561.200.

*Synthesis of 7-[bis(tert-butoxycarbonylmethyl)amino]-4-[({[12-(aminooxy)-1,4,7,10-tetraoxadodecanyl]carbamoyl}oxy)methyl]coumarin 7*

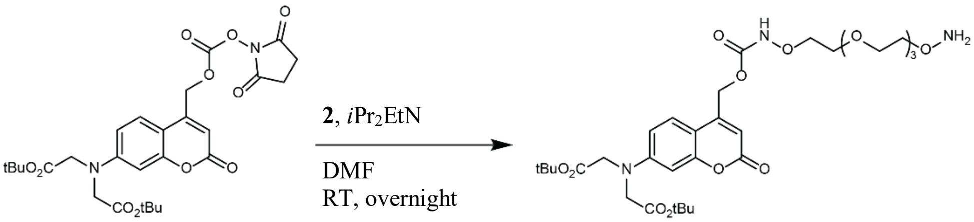

**6** **7**

**3** (1000 mg, 1.8 mmol) and **6** (940 mg, 4.2 mmol) were dissolved in 10 mL of dry *N,N*-dimethylformamide in a flame-dried round bottom flask purged with $N_2$. *N,N*-diisopropylethylamine (470 mg, 3.6 mmol) was added, and the reaction was allowed to continue overnight at room temperature in the dark. Solvent was removed *in vacuo*, and the crude product was dissolved in 1:1 acetonitrile:water and purified by preparatory C18 HPLC using a gradient of 50-100% acetonitrile in water with 0.1% trifluoroacetic acid to yield 490 mg of **7** (0.7 mmol, 41%), a light yellow solid. $^1$H NMR (500 MHz, $CDCl_3$): δ = 1.48 (s, 18 H), 3.56-3.70 (m, 9 H), 3.70-3.74 (t, J = 4.4 Hz, 2 H), 3.81-3.88 (t, J = 3.8 Hz, 2 H), 4.05 (s, 4 H), 4.06-4.09 (m, 2 H), 4.20-4.24 (t, J = 3.7 Hz, 2 H), 5.28 (s, 2 H), 6.19 (s, 1 H), 6.44-6.46 (d, J = 2.6 Hz, 1 H), 6.50-6.55 (dd, J = 2.7 Hz, 9.0 Hz), 7.35-7.39 (d, J = 9.0 Hz, 1 H). MS (ESI-MS): calculated for $C_{31}H_{48}N_3O_{13}$ ($[M+H]^+$): 670.318; found: 670.316.

*Synthesis of 7-[bis(carboxymethyl)amino]-4-[({[12-(aminooxy)-1,4,7,10-tetraoxadodecanyl]carbamoyl}oxy)methyl]coumarin **8***

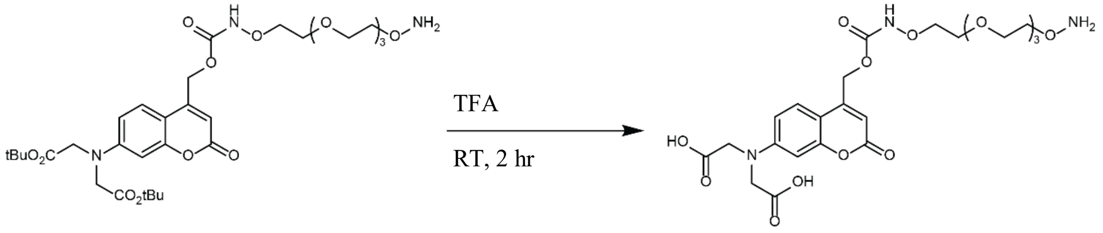

**7** **8**

**7** (442 mg, 0.66 mmol) was dissolved in 10 mL of a 95:2.5:2.5 solution of trifluoroacetic acid:water:triisopropyl silane. After stirring for 2 hours, the solution diluted into cold diethyl ether, then centrifuged for 5 minutes at 5000xg. The solvent phase was decanted, and solids were dried under $N_2$ gas. The pellet was dissolved in 1:9 acetonitrile:water and purified by preparatory C18 HPLC using a gradient of 10-100% acetonitrile in water with 0.1% trifluoroacetic acid to yield 206 mg of **8** (0.37 mmol, 56%), a fluffy light yellow solid. $^1$H NMR (500 MHz, DMSO-$d_6$): 3.65-3.72 (m, 2 H), 3.75-3.81 (m, 2 H) 3.96-4.04 (m, 2 H), 4.25-4.31 (m, 2 H), 4.41 (s, 4 H), 5.40 (s, 2 H), 6.11 (s, 1 H), 6.54-6.59 (d, J = 2.5 Hz, 1 H), 6.68-6.76 (dd, J = 2.9 Hz, 8.9 Hz, 1 H), 7.54-7.65 (d, J = 8.9 Hz, 1 H), 8.01 (s, 1 H), MS (ESI-MS): calculated for $C_{23}H_{32}N_3O_{13}$ ($[M+H]^+$): 558.193; found: 558.194.

*Synthesis of 7-[bis(carboxymethyl)amino]-4-[({[12-[({[(1R,8S,9S)-bicyclo[6.1.0]non-4-yn-9-yl]methoxy}carbonyl)aminoxy]-1,4,7,10-tetraoxadodecanyl]carbamoyl}oxy)methyl]coumarin (DCMAC-HNO-TEG-BCN) **9***

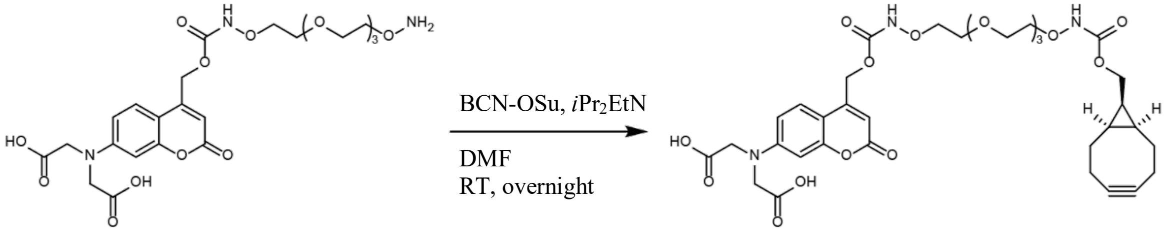

**8** **9**

**8** (53 mg, 0.10 mmol) and (1R,8S,9s)-Bicyclo[6.1.0]non-4-yn-9-ylmethyl *N*-succinimidyl carbonate (BCN-OSu, 41 mg, 0.14 mmol) were dissolved in 1 mL of dry *N,N*-dimethylformamide in a vial purged with $N_2$. *N,N*-diisopropylethylamine (23 mg, 0.18 mmol) was added, and the reaction was allowed to continue overnight at room temperature in the dark. Solvent was removed *in vacuo*. The crude product was dissolved in 9:10 acetonitrile:water and purified by preparatory C18 HPLC using a gradient of 45-100% acetonitrile in water with 0.1% trifluoroacetic acid to yield 44 mg of **9** (DCMAC-HNO-TEG-BCN, 0.06 mmol, 63%), a light yellow gum. $^1$H NMR (500 MHz, $CD_3CN$): δ = 0.90-0.98 (m, 2 H), 1.38-1.65 (m, 3 H), 2.13-2.33 (m, 6 H), 3.56-3.76 (m, 12 H), 3.90-3.95 (m, 2 H), 3.96-4.01 (m, 2 H), 4.17-4.22 (d, J = 8.2 Hz, 2 H), 4.29 (s, 4 H), 5.28 (s, 2 H), 6.15 (s, 1 H), 6.47-6.50 (d, J = 2.2 Hz, 1 H), 6.57-6.62 (d, J = 8.7 Hz, 1 H), 7.47-7.52 (d, J = 8.9 Hz, 1 H), 8.51 (br. s, 1 H), 8.89 (br. s, 1 H). MS (ESI-MS): calculated for $C_{34}H_{44}N_3O_{15}$ ($[M+H]^+$): 734.277; found: 734.278.

**Method S11 Hydrogel synthesis for biomolecule photoimmobilization studies**

$N_3$-PEG-$N_3$ (8 mM final concentration) and DCMAC-HNO-TEG-BCN (100 μM final concentration) in PBS were reacted (30 min, 37 °C). The volume was adjusted with PBS before addition of 10 kDa PEG-tetraBCN (4 mM final concentration). The solution was briefly vortexed and benchtop centrifuged to remove bubbles before pipetting onto a microscope slide double coated with Rain-X. A second RainX-treated slide was placed on top of the droplets separated from the bottom slide by silicon gaskets (0.5 mm thick, McMaster-Carr). The slides were covered in aluminum foil and left at room temperature for 45 minutes to allow gelation. Gels were transferred to a multiwell plate containing PBS and dilute (<10 μM) sodium azide or azide-PEG3-amine ($N_3$-TEG-$N_3$, Lumiprobe) to fully hydrate, effectively "capping" any unreacted BCN while washing out of unbound biomolecules. Gels formed in this manner could be patterned immediately or kept at room temperature in the dark for at least two weeks before patterning. DCMAC's intrinsic fluorescence proved helpful for gel visualization prior to GIZMO patterning.

**Figure S3 Photoimmobilization of EGFP-CHO in shape of Seattle Space Needle**

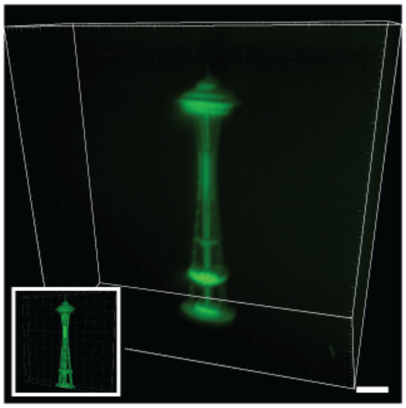

3D patterned immobilization of EGFP-CHO is achieved via photomediated oxime within a DCMAC-HNO-TEG-BCN-modified gel in the shape of the Seattle Space Needle. Scale bar = 50 μm.

**Method S12 Hydrogel synthesis for photodegradation studies**

PEG-tetraBCN ($M_n \approx 20$ kDa, 4.05 mM final concentration) was prereacted with AlexaFluor 488-azide (0.025 mM final concentration, used for gel visualization prior to GIZMO patterning) in PBS. After 1 h, diazide-containing Rubpy (8 mM final concentration) was added in PBS. Gelation was allowed to proceed for 1 h between Rain-X-treated glass slides with silicone rubber spacers (1 mm thick, McMaster-Carr). Following slide separation, gels were equilibrated overnight in PBS prior to use.

Nonphotodegradable control gels were similarly synthesized, substituting $N_3$-TEG-$N_3$ (8 mM) for Rubpy.

**Figure S4 Gel labeling following patterning illustrates extent of network degradation**

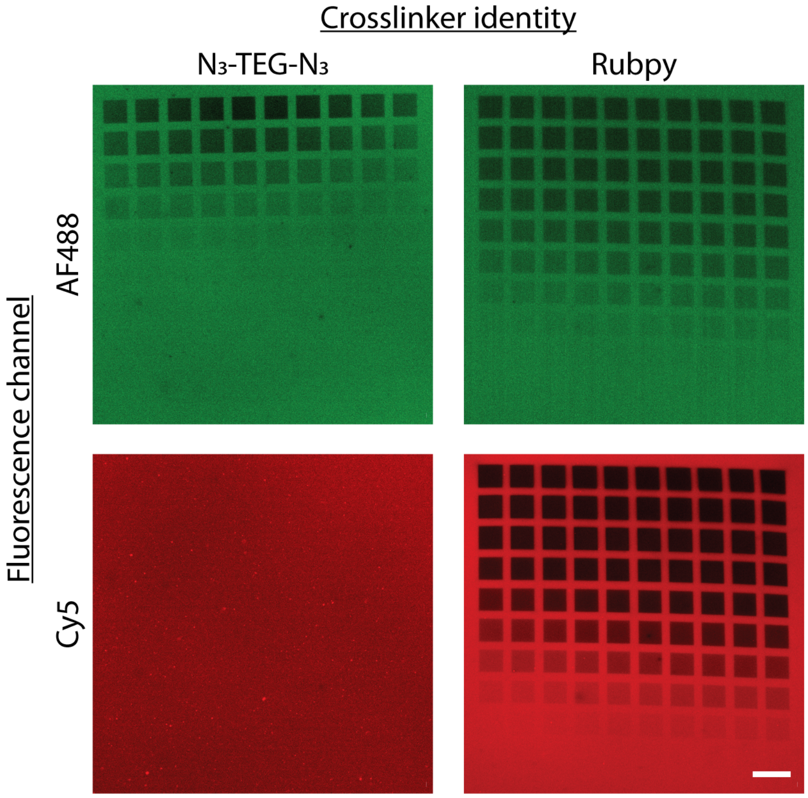

Both photodegradable and control hydrogels (respectively crosslinked with Rubpy and $N_3$-TEG-$N_3$) were cast with AF488 as described in Method S12 prior to GIZMO patterning. Here, gels were exposed to a gridded array of varying power (0% – 100% relative to maximum, moving from bottom-left to top-right of each image). Following light exposure, gels were labeled with Cy5-$N_3$ and then imaged. For the control gel, patterns in the AF488 channel correspond to bleaching; no photoablation was observed in the Cy5 channel. Grayscale gel softening accompanied GIZMO patterning of the Rubpy gels, as indicated by variable Cy5 fluorescence. Scale bar = 50 μm.

**Figure S5 Photodegradation of patent microvessels within gels**

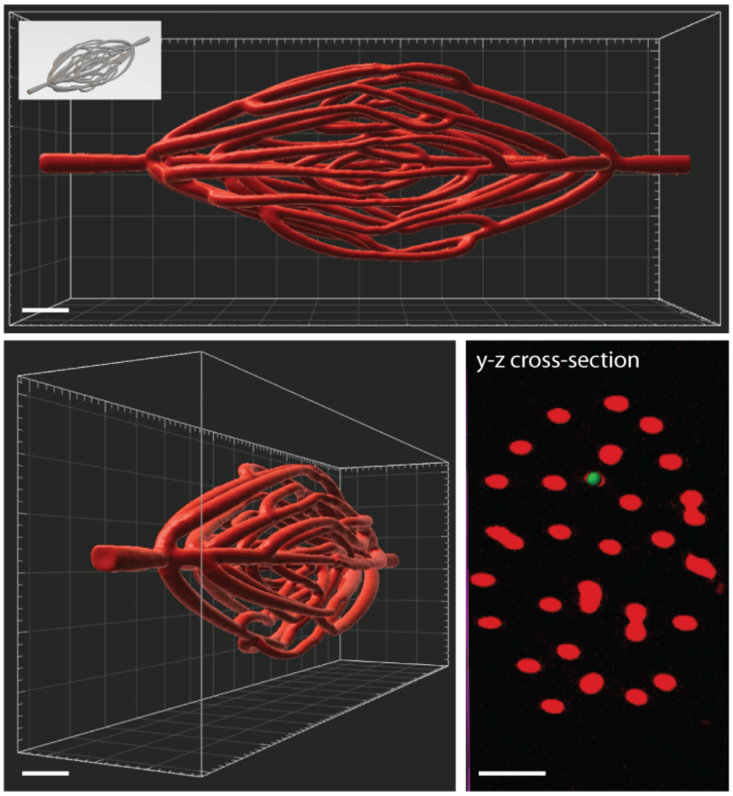

Dendritic vascular network channels computationally grown within an ellipsoidal domain[9] were photodegraded via GIZMO within Rubpy-crosslinked hydrogels. Following photodegradation, gels were soaked in a solution containing fluorescein isothiocyanate-dextran ($M_n$ ~ 2,000 kDa, 1 mg mL$^{-1}$, Sigma Aldrich FD2000S), which was free to diffuse through the patterned voids but not into the intact gel. The fluorescein channel was imaged ~10 hours later (shown above in red), demonstrating high patency and resolution of the patterned voids that matched the inputted structure (inset image). Green circle in y-z cross-section image is shown at 10 μm diameter, reflecting a typical native capillary size. Scale bars = 50 μm.

### Method S13 Clearing, 3D staining, and imaging of whole mouse brain

Clearing, 3D staining, and imaging of whole mouse brain samples were performed with CUBIC clearing and CUBIC-HistoVision (CUBIC-HV) staining protocols[10,11] as we have reported previously[12]. An updated CUBIC-HV staining protocol was used (HV1.1, commercialized by CUBICStars Co. and Tokyo Chemical Industry (TCI), TCI no. C3709, no. C3708). In brief, the PFA-fixed whole mouse brains were treated with CUBIC-L for 4 days at 37 °C, washed with PBS, stained with SYTOX-G (1/2500) in CUBIC-HV nuclear-staining buffer (included in TCI no. C3709) for 5 days at 37 °C, and then washed and immersed in 50% and 100% CUBIC-R + for 1 day and 3 days, respectively, at room temperature. We used CUBIC-R + (N) (45 wt% of antipyrine, 30 wt% of nicotinamide (TCI no. N0078), and 0.5% (v/w) N-butyldiethanolamine, adjusted to pH ~10).

For the whole mouse brain immunostained with anti-α-SMA (Sigma, no. A5228) antibodies, the brain was subjected to CUBIC-HV immunostaining. The brain was first treated with 3 mg $mL^{-1}$ hyaluronidase in CAPSO buffer (pH 10) for 24 hours at 37 °C. After washing with hyaluronidase wash buffer (50 mM carbonate buffer, 0.1% (v/v) Triton X-100, 5% (v/v) methanol, and 0.05% $NaN_3$) and HEPES-TSC buffer (10 mM HEPES buffer, pH 7.5, 10% (v/v) Triton X-100, 200 mM NaCl, 0.5% (w/v) casein, and 0.05% $NaN_3$), the brain was immersed in 500 μL of HEPES-TSC buffer containing a primary antibody (6 μg for anti-α-SMA), a secondary Fab fragment (FabuLight, Jackson immunolab, Alexa Fluor 594 goat anti-mouse IgG1 no. 115-587-185, Alexa Fluor 594 goat anti-rabbit IgG no. 111-587-008, Alexa Fluor 594 goat anti-mouse IgG2a no. 115-587-186, 1:0.75 of weight ratio), and 3D immunostaining additive (1×) (included in TCI no. C3708). Then, the sample was incubated with gentle shaking for 10 days at 32 °C. After staining, the sample was additionally incubated in the same buffer for 1 day at 4 °C to stabilize the Fab binding. Then, the sample was washed and post-fixed according to the protocol of CUBIC-HV immunostaining kit (TCI no. C3708) before being index-matched with CUBIC-R+. The cleared sample was embedded in CUBIC-R-agarose for imaging and storage[13]. For CUBIC-cleared tissues, which are less rigid than solvent-cleared tissues, agarose embedding helps to stabilize the tissues during imaging. This facilitates alignment of the resulting datasets across tiles and channels, as well as co-registration of ODO and NODO datasets. The animal experimental procedures and housing conditions of the animals were approved by the Animal Care and Use Committees of the Graduate School of Medicine of the University of Tokyo.

The entire intact brain was imaged at a rate of ~300 – 400 $mm^3$ $hr^{-1}$ with near-isotropic resolution (*xyz* resolution of 4.41 ± 0.83, 4.09 ± 1.07, and 5.48 ± 1.08 μm) using the mesoscopic ODO path of the hybrid open-top light-sheet (OTLS) system, which clearly resolves the vasculature in all three dimensions.

**Method S14 3D mouse brain blood vessel segmentation**

We developed a custom Python-based workflow for 3D vessel segmentation in light-sheet images of mouse brain samples stained with CD31 that highlights the blood vessel walls. The segmentation process mainly consisted of two steps.

Firstly, we segmented the vessel walls using localized multi-thresholding followed by hysteresis thresholding[14]. Specifically, a cubic box was used to scan through the entire 3D data with overlap between each adjacent step. At each location of the sliding box, we applied Otsu multi-thresholding to the box to identify the foreground region (i.e., vessel walls), the background region (i.e., non-vessel walls), and the "gray-area" in between with vague classification, based on the distribution of intensities within the box. Hysteresis thresholding was then applied to classify any "gray-area" that was connected to the foreground as foreground, while the rest of the "gray-area" was classified as background. This approach allowed for more complete vessel wall segmentation, even in cases of non-uniform CD31 staining or vessels with small cuts.

Secondly, we segmented the inner vessel region using a slice-by-slice contour-filling routine[15]. This routine involved filling in the regions enclosed by the vessel wall mask in every 2D slice of the 3D data. The resulting mask of the inner vessel region was then combined with the mask of vessel walls to create a total mask for blood vessels.

Relevant code is available here:
https://github.com/WeisiX/3D_vessel_seg/blob/main/cubic01_4900ROI2mm_stitch_3Dmultiotsu_final.py

**Figure S6 Grayscale softening of Rubpy-crosslinked hydrogels via GIZMO**

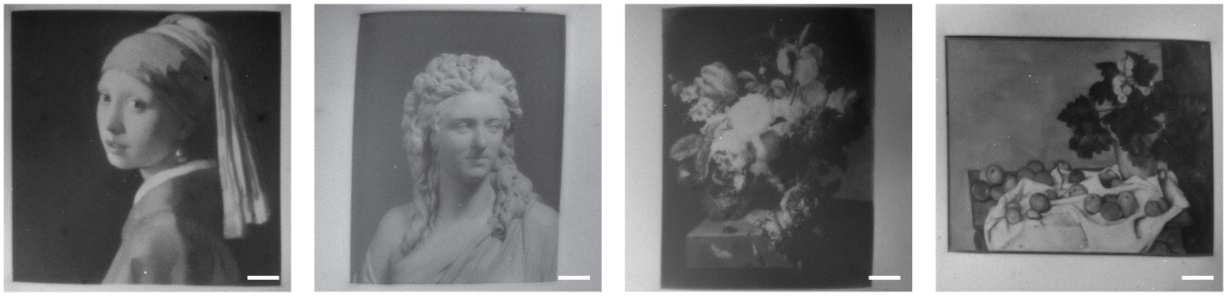

Stiffness-patterned gels were created in the form of Vermeer's "Girl with a Pearl Earring", a photograph of Pajou's statue entitled "Madame de Wailly, née Adélaïde-Flore Belleville", Faes's "Flowers in a Stone Vase", and Cézanne's "Still Life with Apples and a Pot of Primroses". Following GIZMO-mediated photosoftening, gels were labeled with Cy5-azide prior to fluorescent imaging. Here, higher fluorescence (white) indicates the natively intact hydrogel, while black indicates regions that are fully photodegraded; grayscale values in between correspond to partially degraded materials of intermediate stiffnesses. Scale bars = 50 µm.

**Movie S1 Peptide-patterned hydrogel "flipbook" via grayscale biomolecule photorelease**
Video comparing inputted drone footage (left) with the GIZMO-patterned biomolecule "flipbook" (right). Greyscale 3D flipbook comprised of fluorescent RGDS peptide is visualized moving from the top to the bottom of its z-stack (300 μm x 300 μm x 1.2 mm total size), stepping through each of the 200 distinct planar images at a frame rate matching the input video.

**Movie S2 Protein-patterned hydrogel "flipbook" via grayscale biomolecule photoimmobilization**
Video comparing inputted video footage (Walt Disney's short film "The Mad Doctor", public domain, top right inset) with the GIZMO-patterned protein "flipbook". Greyscale 3D flipbook comprised of mGL-CHO is visualized moving from the top to the bottom of its z-stack (400 μm x 300 μm x 600 μm total size), stepping through each of the 200 distinct planar images at a frame rate matching the input video.

**Movie S3 Anatomically matched photodegraded vessels in biomaterials**
Video showcasing the full mouse brain dataset obtained through tissue clearing, immunohistochemical labeling, and OTLS imaging, with the 2 mm x 2 mm x 2 mm subvolume-of-interest is highlighted in cyan, the results of vessel segmentation in purple, and tiled GIZMO-photodegraded voids within hydrogel in green.

**Movie S4 Flythrough of GIZMO-patterned microvessel network**
Surface renderings of tiled GIZMO-photodegraded voids throughout gel following native mouse brain microvasculature pattern.

**Movie S5 Stiffness-patterned hydrogel "flipbook" via grayscale network photodegradation**
Video comparing inputted video footage (Popeye short film "A Date to Skate", public domain, top right inset) with the GIZMO-patterned stiffness "flipbook". Greyscale 3D flipbook is visualized moving from the top to the bottom of its z-stack (450 μm x 350 μm x 650 μm total size), stepping through each of the 300 distinct planar images at a frame rate matching the input video.

### References

1. DeForest, C. A. & Tirrell, D. A. A photoreversible protein-patterning approach for guiding stem cell fate in three-dimensional gels. *Nature Materials* **14,** 523–531 (2015).
2. Badeau, B. A., Comerford, M. P., Arakawa, C. K., Shadish, J. A. & DeForest, C. A. Engineered modular biomaterial logic gates for environmentally triggered therapeutic delivery. *Nature Chemistry* **10,** 251–258 (2018).
3. Rapp, T. L. & DeForest, C. A. Tricolor visible wavelength-selective photodegradable hydrogel biomaterials. *Nat Commun* **14,** 5250 (2023).
4. Shadish, J. A., Benuska, G. M. & DeForest, C. A. Bioactive site-specifically modified proteins for 4D patterning of gel biomaterials. *Nature Materials* **18,** 1005–1014 (2019).
5. Munoz-Robles, B. G. & DeForest, C. A. Irreversible light-activated SpyLigation mediates split protein assembly in 4D. *Nature Protocols* (2024). doi:10.1038/s41596-023-00938-0
6. Broguiere, N., Lüchtefeld, I., Trachsel, L., Mazunin, D., Rizzo, R., Bode, J. W., Lutolf, M. P. & Zenobi-Wong, M. Morphogenesis Guided by 3D Patterning of Growth Factors in Biological Matrices. *Advanced Materials* **32,** 1908299 (2020).
7. Gieß, M., Muñoz-López, Á., Buchmuller, B., Kubik, G. & Summerer, D. Programmable Protein–DNA Cross-Linking for the Direct Capture and Quantification of 5-Formylcytosine. *Journal of the American Chemical Society* **141,** 9453–9457 (2019).
8. Broguiere, N., Lüchtefeld, I., Trachsel, L., Mazunin, D., Rizzo, R., Bode, J. W., Lutolf, M. P. & Zenobi-Wong, M. Morphogenesis Guided by 3D Patterning of Growth Factors in Biological Matrices. *Advanced Materials* **32,** 1908299 (2020).
9. Kinstlinger, I. S., Saxton, S. H., Calderon, G. A., Ruiz, K. V., Yalacki, D. R., Deme, P. R., Rosenkrantz, J. E., Louis-Rosenberg, J. D., Johansson, F., Janson, K. D., Sazer, D. W., Panchavati, S. S., Bissig, K.-D., Stevens, K. R. & Miller, J. S. Generation of model tissues with dendritic vascular networks via sacrificial laser-sintered carbohydrate templates. *Nature Biomedical Engineering* **4,** 916–932 (2020).
10. Tainaka, K., Murakami, T. C., Susaki, E. A., Shimizu, C., Saito, R., Takahashi, K., Hayashi-Takagi, A., Sekiya, H., Arima, Y., Nojima, S., Ikemura, M., Ushiku, T., Shimizu, Y., Murakami, M., Tanaka, K. F., Iino, M., Kasai, H., Sasaoka, T., Kobayashi, K., Miyazono, K., Morii, E., Isa, T., Fukayama, M., Kakita, A. & Ueda, H. R. Chemical Landscape for Tissue Clearing Based on Hydrophilic Reagents. *Cell Reports* **24,** 2196-2210.e9 (2018).
11. Susaki, E. A., Shimizu, C., Kuno, A., Tainaka, K., Li, X., Nishi, K., Morishima, K., Ono, H., Ode, K. L., Saeki, Y., Miyamichi, K., Isa, K., Yokoyama, C., Kitaura, H., Ikemura, M., Ushiku, T., Shimizu, Y., Saito, T., Saido, T. C., Fukayama, M., Onoe, H., Touhara, K., Isa, T., Kakita, A., Shibayama, M. & Ueda, H. R. Versatile whole-organ/body staining and imaging based on electrolyte-gel properties of biological tissues. *Nat Commun* **11,** 1982 (2020).
12. Glaser, A. K., Bishop, K. W., Barner, L. A., Susaki, E. A., Kubota, S. I., Gao, G., Serafin, R. B., Balaram, P., Turschak, E., Nicovich, P. R., Lai, H., Lucas, L. A. G., Yi, Y., Nichols, E. K., Huang, H., Reder, N. P., Wilson, J. J., Sivakumar, R., Shamskhou, E., Stoltzfus, C. R., Wei, X., Hempton, A. K., Pende, M., Murawala, P., Dodt, H.-U., Imaizumi, T., Shendure, J., Beliveau, B. J., Gerner, M. Y., Xin, L., Zhao, H., True, L. D., Reid, R. C., Chandrashekar, J., Ueda, H. R., Svoboda, K. & Liu, J. T. C. A hybrid open-top light-sheet microscope for versatile multi-scale imaging of cleared tissues. *Nature Methods* **19,** 613–619 (2022).

13. Matsumoto, K., Mitani, T. T., Horiguchi, S. A., Kaneshiro, J., Murakami, T. C., Mano, T., Fujishima, H., Konno, A., Watanabe, T. M., Hirai, H. & Ueda, H. R. Advanced CUBIC tissue clearing for whole-organ cell profiling. *Nat Protoc* **14,** 3506–3537 (2019).
14. Bukenya, F., Awwad, A., Duan, J., Ehling, J., Faas, H. & Bai, L. Three-dimensional Segmentation of Blood Vessels from Intensity In-homogeneous Medical Images. in *2018 IEEE Symposium Series on Computational Intelligence (SSCI)* 1508–1514 (IEEE, 2018). doi:10.1109/SSCI.2018.8628675
15. Xie, W., Reder, N. P., Koyuncu, C., Leo, P., Hawley, S., Huang, H., Mao, C., Postupna, N., Kang, S., Serafin, R., Gao, G., Han, Q., Bishop, K. W., Barner, L. A., Fu, P., Wright, J. L., Keene, C. D., Vaughan, J. C., Janowczyk, A., Glaser, A. K., Madabhushi, A., True, L. D. & Liu, J. T. C. Prostate Cancer Risk Stratification via Nondestructive 3D Pathology with Deep Learning–Assisted Gland Analysis. *Cancer Research* **82,** 334–345 (2022).
